# Representations in the hippocampal-entorhinal system emerge from learning sensory predictions

**DOI:** 10.1101/2025.10.03.680189

**Authors:** Diego Gomez, Michael Bowling, J. Quinn Lee, Marlos C. Machado

## Abstract

The hippocampal formation and adjacent parahippocampal areas are central to intelligent behaviour such as memory and navigation. Understanding how systems in the brain generate structured representations from experience remains a fundamental goal in neuroscience. A central open question is whether a single computational principle can account for the diverse neural responses observed across the hippocampal-entorhinal circuit. Existing models often rely on hand-crafted features or specialized learning mechanisms unrelated to sensory observations, and typically express representations of only a small subset of known cell types. Further, representations learned in such models are often not empirically evaluated against neural representations observed in the navigating brain. Here, we introduce a neurobiologically-inspired and robust computational model in which diverse cell types emerge from a single learning objective with minimal hand-engineered assumptions. Our model applies contrastive graph representation learning to transitions between high-dimensional visual observations, constructing a metric space in which temporally adjacent sensory observations are mapped to nearby states. Inspired by the anatomical information flow of the hippocampal-entorhinal system, and anchored in output representations based on neural coding in the entorhinal cortex, the model gives rise to activity resembling place cells, grid cells, boundary vector cells, band cells, corner cells, and conjunctive cells among others. Across varied environments and sensory streams, the framework captures not only diverse neural response patterns but also the functional dependencies between them, mirroring the proposed sequential representational structure observed in the hippocampal-entorhinal system. Crucially, place-cell-like features of the model quantitatively reproduce remapping dynamics observed in CA1 of freely moving animals, and afford theoretical explanatory power of existing neurobiologically-informed models. This work thus offers a unified computational model of spatial coding in the hippocampal-entorhinal system and a testable framework for generating mechanistic hypotheses *in silico*, to be evaluated *in vivo*.

---

Understanding hippocampal neural representations and their role in intelligent behaviour, including memory, navigation, and planning has been a long-standing goal since the discovery of place cells and grid cells (*1*). A key remaining question is whether there is a single computational principle that might account for the characteristic representations across subregions of the hippocampal-entorhinal system. A growing number of computational models suggest that the hippocampus functions as a predictive machine, encoding sensory and internal signals in an abstract state-space to facilitate the prediction and control of future experiences (*2–13*). However, the relationship between computation in recent models and their analogous neurobiological systems remains poorly understood, with competing models proposing different mechanisms for the interdependence of cell types (*2, 14–18*), the specific class of predictive representations (*4, 10, 19*), and the underlying learning algorithms (*18, 20*).

Most models of hippocampal spatial coding assume that animals either have access to explicit spatial variables, such as Euclidean coordinates or distances to external boundaries, or that each state is specified by an abstract symbolic identifier, such as a one-hot code or unique state index (*15–17, 21, 22*). These models typically describe neural responses as analytic functions of these variables, or as hierarchical combinations. While they reproduce key neural response patterns and explain relationships between specific cell types, such models rely on biologically implausible assumptions about observations available to the animal, and in most cases provide no account of how these representations could be learned from experience (*8*). More recent models have shown that grid cells and other hippocampal responses—like replay (*10*)—can emerge through learning, generally via some form of path integration in artificial neural networks (*3–6, 23*) or variants of the successor representation (SR) (*19,24,25*). Yet, existing path integration models assume pre-existing localization (e.g., head-direction cells and place cells) (*3*) or self-motion (*5*) (e.g., translational and angular velocities) systems and do not account for known sequential dependencies between cell types (*15, 16, 26, 27*). SR-based models, by contrast, typically rely on discrete state spaces (*19*) or require strong assumptions about the inputs (*24, 25*). Furthermore, these models offer limited or likely implausible explanations of how place and grid cells relate, often invoking computations such as eigen-decomposition or outer product combinations that available evidence does not suggest is plausible in the brain (*19, 24*).

## Learning algorithm

In our model, we learn a representation that is comprised of a set of features, where each feature can be seen as representing a different cell. Inspired by the role the dentate gyrus is believed to have in orthogonalization (*28,29*), these features are designed to be orthogonal to each other, and, together, they are learned to represent observations that are close in time as nearby states. Specifically, we instantiate this model by training a neural network to optimize a formulation of this objective, which was shown (*30*) to produce an approximation of the first eigenvectors of the graph Laplacian—120 in this case—induced by observed sensory transitions, ordered by eigenvalue. These eigenvectors embed observations into a metric space in which the distance reflects the expected transition time; that is, temporally adjacent states are placed nearby. In a simplified setting, under a symmetric environment and a uniform random policy, these Laplacian eigenvectors are equivalent to those of the successor representation (*30, 31*).

To learn these eigenvectors, we optimize the augmented Lagrangian Laplacian objective (ALLO) (*30*):

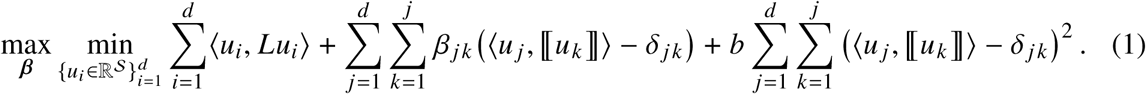

The first term encourages temporal smoothness by aligning the *i*-th component of the representation, *u*_*i*_, with the corresponding eigenvector of the subsequent observation, *Lu*_*i*_. The remaining terms orthogonally constrain components captured by the augmented Lagrangian formulation, where δ _*j k*_ denotes the Kronecker delta (see (*32*) for details). Together, these terms ensure that each component learns a temporally coherent and spatially distinct embedding dimension. Notably, the orthogonalization can be viewed as a computational analogue, in terms of objective function rather than architectural design, to the sparsification and decorrelation believed to occur through combined inhibition and expansion in connectivity and population size from the entorhinal cortex to the dentate gyrus (*28, 29*).

In this paper, we focused on training a deep neural network on data generated from simulated rodent trajectories in a 3D open arena designed to mimic conditions from recent empirical studies (*33*) (Fig. 1A). Each trajectory was represented as a sequence of egocentric pixel-based images of size 60× 80×3 (Fig. 1B). During training, the network received batches of observation triples composed of consecutive and randomly sampled (non-adjacent) frames (see (*32*) for details). The architecture consisted of a three-layer convolutional encoder followed by a four-layer fully connected module with ReLU activations, enforcing non-negativity (Fig. 1C). Importantly, the network received no allocentric inputs such as position, velocity, or head direction, and no architectural regularization (e.g., dropout, batch normalization) was used, in contrast to prior models (*3*). Note that although we focused in our experiments on using optical flow as input, our model is quite general as it can receive any type of sensory observation. Other types of sensory observation might require adjustments in the first layers of the neural network, as convolutional layers are commonly used for image processing. Still, the objective function we optimize would remain unchanged.

**Figure 1:**
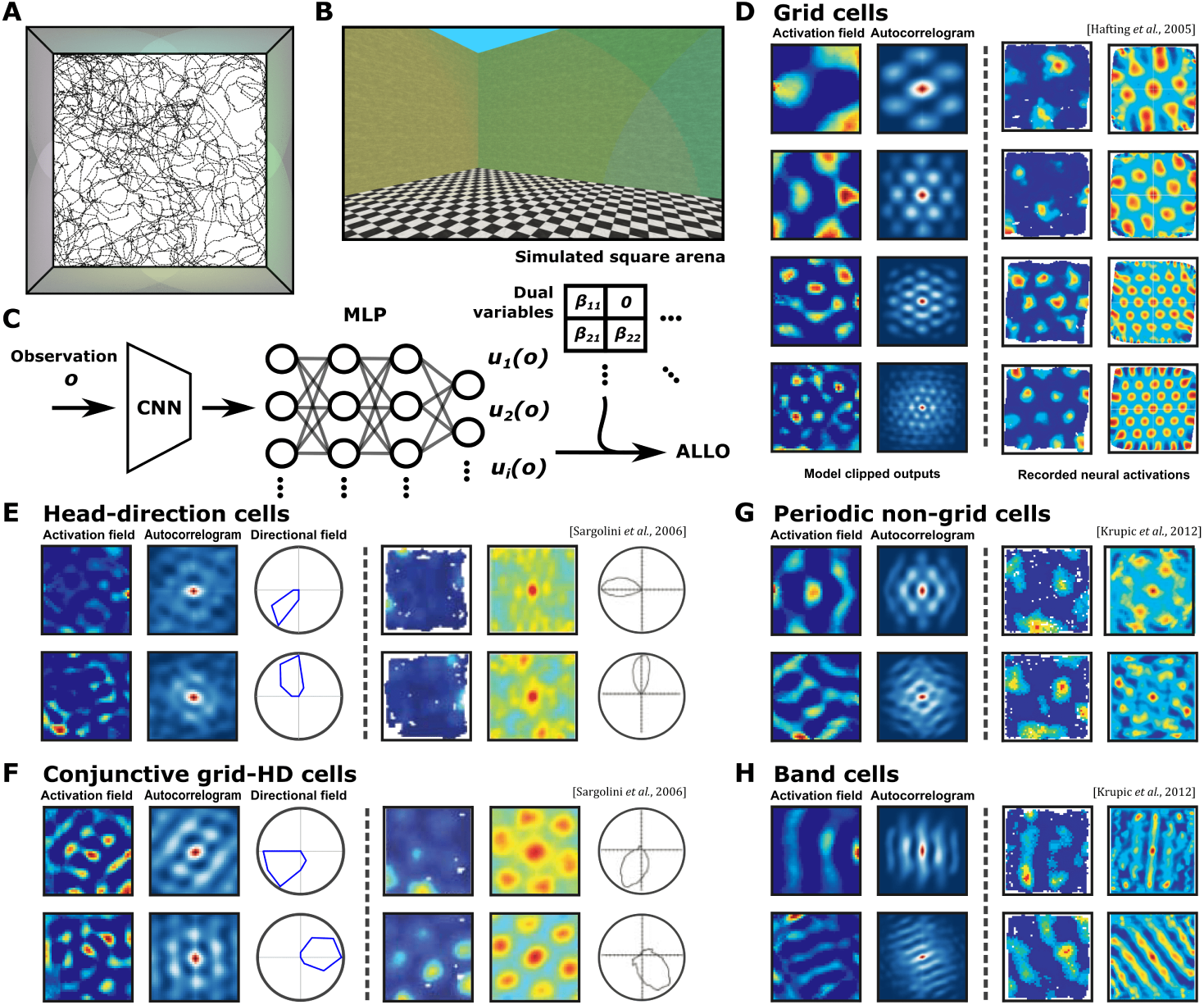
Model eigenvectors exhibit spatial tuning matching neurons in the medial entorhinal cortex, pre- and parasubiculum. (**A**) Schematic of the square open arena used to simulate rodent trajectories (dotted line). Colored lighting in the MiniWorld environment to create polarizing visual stimuli at each wall. (**B**) Example of a high-resolution egocentric observation; input to the model consists of RGB images of size 60 × 80 × 3. (**C**) Architecture of the eigenvector deep neural network, composed of a convolutional encoder and a fully connected module. Observations *o* are fed into the network and the resulting outputs *u*_*i*_ (*o*) are used to calculate the ALLO objective together with the dual variables β_*i j*_. (**D–H**) Comparison between model features (left columns) and firing patterns observed in rodent studies (right columns). For each model feature, the spatial activation map, autocorrelogram (second-left column), and polar plot are shown (third-left column) where applicable. (**D**) Grid cells, arranged by eigenvalue; grid spacing in the model decreases monotonically mirroring the the entorhinal cortex behavior. (**E**) Head-direction cells: spatially diffuse, but strongly direction-tuned responses (third column shows angular tuning). (**F**) Conjunctive cells exhibit both spatial periodicity and head-direction tuning (third column shows angular tuning). (**G–H**) Periodic but non-grid-like cells (**G**), including band cells (**H**).

## Results

### Parahippocampal-, pre- and parasubicular-type feature emergence

A wide range of parahippocampal and pre- and parasubicular neurons exhibit spatial tuning, including grid cells, band cells, and conjunctive cells among others (*34, 35*). These responses are often periodic and can be described using a small number of Fourier components (*35*). Among them, grid cells are perhaps the most widely studied: their firing fields exhibit hexagonal symmetry across the environment, effectively providing the animal with a localization scaffold (*34, 36*). Prior work has shown that predictive representations, such as the eigenvectors of the successor representation (*37*) or of the graph Laplacian (due to their equivalence in some settings (*31*)), can give rise to grid-like responses (*19, 24*), but these works have typically focused on successor features with place-like activations and grid cells alone. In contrast, we hypothesize that the broader set of eigenvectors of the graph Laplacian can account for a wider diversity of spatially modulated cells across parahippocampal and pre- and parasubicular regions, and not only place- or grid-like features.

After training the neural network to optimize the ALLO objective and, as a result, to approximate the eigenvectors of the graph Laplacian (see (*32*) for details), we evaluated the activation of each output unit across positions in a regular spatial grid to construct spatial activation maps, analogous to the tuning curves used to categorize spatially-tuned neurons in experimental studies. To illustrate the resemblance of our learned features to observed spatial cell types in parahippocampal and pre- and parasubicular regions, we first gathered representative examples of spatially tuned cells from the literature and compared them to the patterns produced by our model across environments with different topologies (see fig. S1).

We first considered grid cells (Fig. 1D), and computed the gridness score of each eigenvector from its autocorrelogram (see (*32*) for details), finding that 24.4% were classified as grid cells, which is in keeping with the proportions reported experimentally in the medial entorhinal cortex and the pre- and parasubiculum (*34, 35*). The autocorrelograms of grid cells exhibit characteristic spacing between the central peak and six surrounding peaks arranged in a hexagonal lattice. This spacing is known to vary inversely with the neuron’s anatomical distance from the postrhinal cortex, which is anatomically adjacent to the entorhinal cortex (*36, 38*). This phenomenon is partially reflected in our model: the spacing between peaks varies systematically with the eigenvalue index (see fig. S2), and this trend is consistent across environment topologies, including square, circular, hexagonal, and trapezoidal arenas (see Fig. 1D, and fig. S3). However, unlike some earlier studies (*39*), we do not observe clear evidence for discretized modules in the distribution of spacings, possibly because of the use of a highly uniform behavior policy to generate trajectories or the low number of eigenvectors used (see fig. S4).

Head-direction cells and conjunctive cells represent another major class of spatially tuned neurons in the parahippocampal-subicular region (in addition to the anterior dorsal thalamic nucleus). Namely, head-direction cells exhibit firing that is selective to head orientation, while conjunctive cells also show grid-like spatial responses that depend on head direction (*34*). We evaluated the directional tuning of the model’s eigenvectors by computing the lengths of their mean weighted Rayleigh vectors (see (*32*) for details). Of the 120 eigenvectors, 46.7% were classified as head-direction cells (Fig. 1E). Among them, 23.2% (and 44.8% of all grid cells) also expressed grid-like modulation and met the criteria for conjunctive cells (Fig. 1F). These proportions are consistent with values reported for layers II, III, V, and VI of the medial entorhinal cortex (49.0% head-direction cells, and 24.1% of those, conjunctive). However, the absolute Rayleigh vector lengths were low, with a maximum value of 0.17, just below the noise threshold of 0.19 typically used to identify head direction cells in experimental literature (*34*).

Border and center-distance cells have also been documented in the same brain regions (*40–43*). Such neurons typically exhibit activations aligned with environmental boundaries, spanning a large fraction of some axes and peaking near walls or at the center of the arena. In our model, the first five eigenvectors showed similar spatial preferences, with some peaking near the center (see fig. S5B) and others exhibiting strong border-related activity (see fig. S5C). Notably, the proportion of such cells in the literature is relatively low, typically ranging from 4% to 12% (*40, 43, 44*), which aligns with the small fraction of model units (3.8%) exhibiting comparable border-related responses (see (*32*) for details).

Next, we analyzed the periodicity of the eigenvectors using their Fourier spectrograms and corresponding polar plots. Following the criteria in (*35*), we classified model neurons as spatially periodic if their strongest Fourier component exceeded the 95th percentile of a null distribution obtained by spatially shuffling the data (see (*32*) for details). Using this criterion, 89.9% of the model’s units were classified as spatially periodic, including 26.2% as grid cells and 56.5% as head-direction cells. The Fourier spectrograms of grid cells typically exhibited three peaks corresponding to different Fourier modes, whereas the remaining spatially periodic but non-grid cells (Fig. 1G) displayed more uniform frequency spectra (see fig. S6). Among those with one, two, or four dominant components, we observed spatial firing fields that resembled band (Fig. 1H) and square patterned activity (see fig. S7), both of which are consistent with cell types recorded in rodents. Notably, these patterns emerged not only in square environments but also in circular arenas (see fig. S1).

### Emergent representations of the hippocampal-entorhinal system

That our model produces a range of cell types recorded in the parahippocampal-subicular region and in a similar proportion suggests these diverse cell types could have a shared computational role, and potentially a common underlying mechanism behind their emergence. This aligns with earlier proposals that spatial responses in this region could be explained from combinations of band patterns, each corresponding to individual Fourier components (Fig. 1G) (*35*). Given that the outputs of our model neurons are linear combinations of features computed in earlier layers, we hypothesized that intermediate activations may also exhibit spatial tuning patterns observed in rodents, including band-like responses in earlier layers. This idea draws inspiration from findings in the visual system, where successive layers of deep convolutional neural networks trained for object recognition have been shown to mirror neural activity across multiple visual areas in primates and humans, including V1, V4, and the inferior temporal cortex (*45*).

Indeed, the output activations of each layer in the fully connected module resembled the firing patterns of cell types found in distinct subregions of the hippocampal formation (Fig. 2A). Specifically, the first and second layers comprised neurons that activated broadly across the arena, often with preferences for one or more boundary-driven patterns that mirror the activity of boundary vector cells (BVCs) and corner cells in the subiculum (*15,46*). In our model, 38.79% and 27.31% of neurons in the first and second layers, respectively, matched BVC-like profiles, while 12.93% and 10.92% resembled corner cells (Figs. 2B,C). These proportions align closely with those reported in adult rats navigating standard square environments (34–36% for BVCs and 4–11% for corner cells) (*44, 46*). Additionally, the directional selectivity of identified BVCs in our model, measured as the ratio between maximum and average polar activation, was 2.12 ± 0.72 in the first layer and 1.88 ± 0.41 in the second. These values are consistent with empirical measurements (1.56 ± 0.05) and are substantially lower than those of head-direction cells (6.91±0.05), and not higher than place cells (2.87 ± 0.20) (*47, 48*). In the third layer, neurons exhibited unimodal and spatially localized responses that, in aggregate, tiled the full arena, closely resembling hippocampal place cells (*49,50*), with a directional selectivity of 2.76 ± 0.84 (Fig. 2D). The final (fourth) layer reproduced periodic responses, including grid-like and band-like activations, matching spatial cell types observed in the medial entorhinal cortex (Fig. 2E).

**Figure 2:**
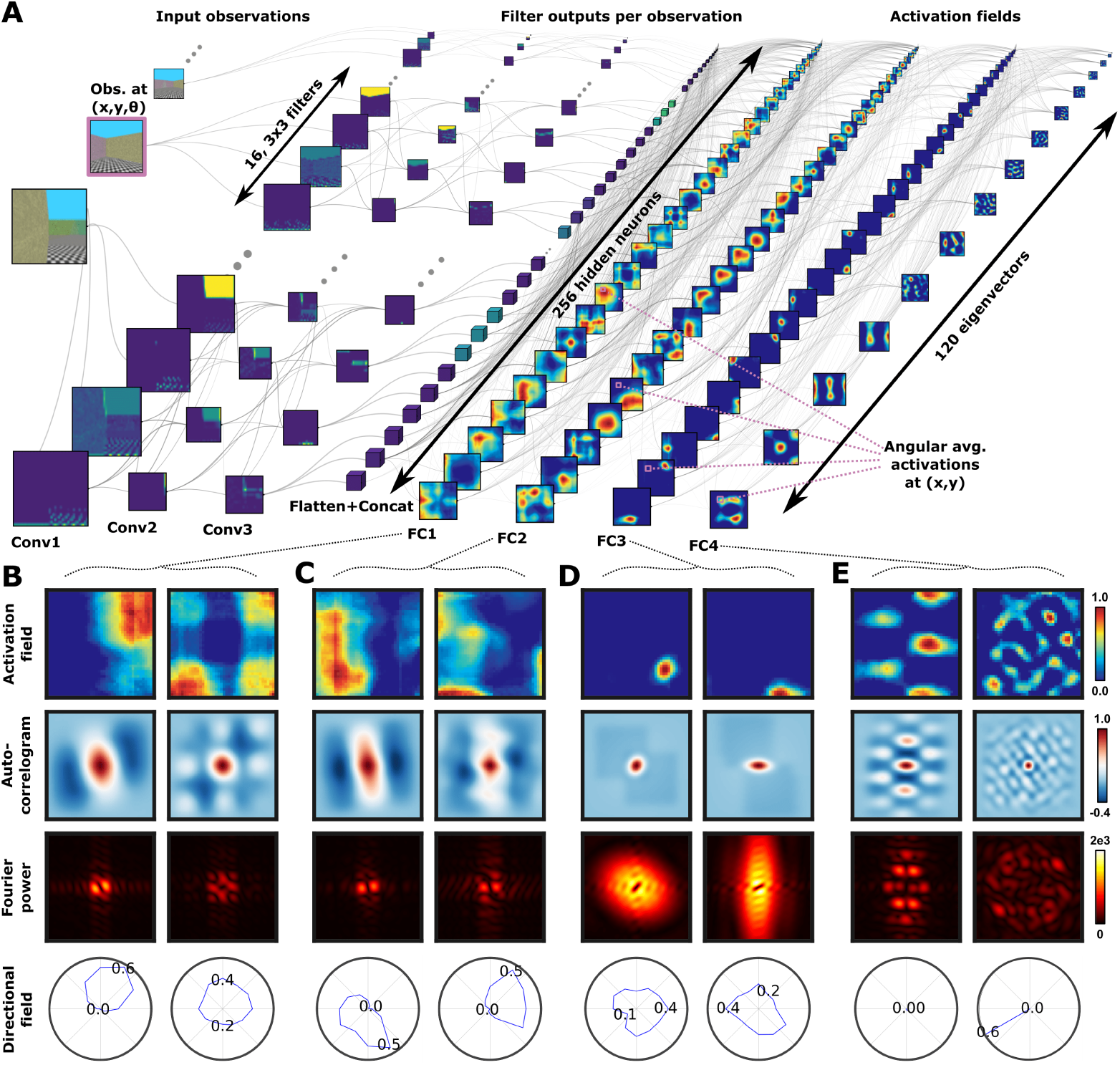
Emergent representations in deep neural networks trained with ALLO express neurobiologically plausible sequential organization. (**A**) Information flow in the model architecture. Each observation, which corresponds to a unique triple of coordinates (*x*, *y*, θ), is processed by a sequence of three convolutional layers, each with 16 filters of kernel size 3 × 3. The result is flattened and concatenated and then fed into the fully connected layers (**FC1**-**FC4**). The spatial activation map in each fully connected node represents all the possible average activations for all possible (*x*, *y*) location pairs. (**B-E**) Learned representations in each of the four MLP layers. For each layer, we show: (first row) example spatial activation maps of representative units; (second row) spatial autocorrelograms; (third row) Fourier spectrograms; and (fourth row) average polar activation profiles. (**B-C**) Border and corner cells in first and second layers. (**D**) Place cells in third layer. (**E**) Grid cell (left) and conjunctive (head direction and spatially-periodic non-grid) cell (right) in final (fourth) layer.

Taken together, these findings suggest that learning to map temporally adjacent sensory observations to nearby locations, when combined with orthogonality constraints, could drive the emergence of specialized cell types observed across the hippocampal-entorhinal system. This includes the subiculum, CA3/CA1, and medial entorhinal cortex, forming a representational loop consistent with known anatomical and functional circuits in the mammalian brain. These results support earlier proposals that place cells may arise from combinations of boundary-sensitive units (*21*), or that they approximate the successor features of BVC activations (*24*).

### Remapping representations aligned with empirical observations in freely behaving rodents

When environments change, place cells adapt their tuning to new locations, a phenomenon known as remapping (*24, 33, 41*). This process offers a useful probe into the representational structure of place cells and serves as a benchmark for model evaluation (*33*). We asses whether the activations in the model’s third fully connected layer, which resemble place cells, also remap similarly to recent observations in CA1 of freely behaving mice (4 males, 3 females), as reported in (*33*).

To this end, we trained the eigenvector model in 10 different geometries, obtained by partitioning a square open arena into a 3 × 3 grid and blocking parts of the space with short walls (Fig. 3A). We simulated observations corresponding to real rodent trajectories and recorded the activations from the third MLP layer (Fig. 3C).

**Figure 3:**
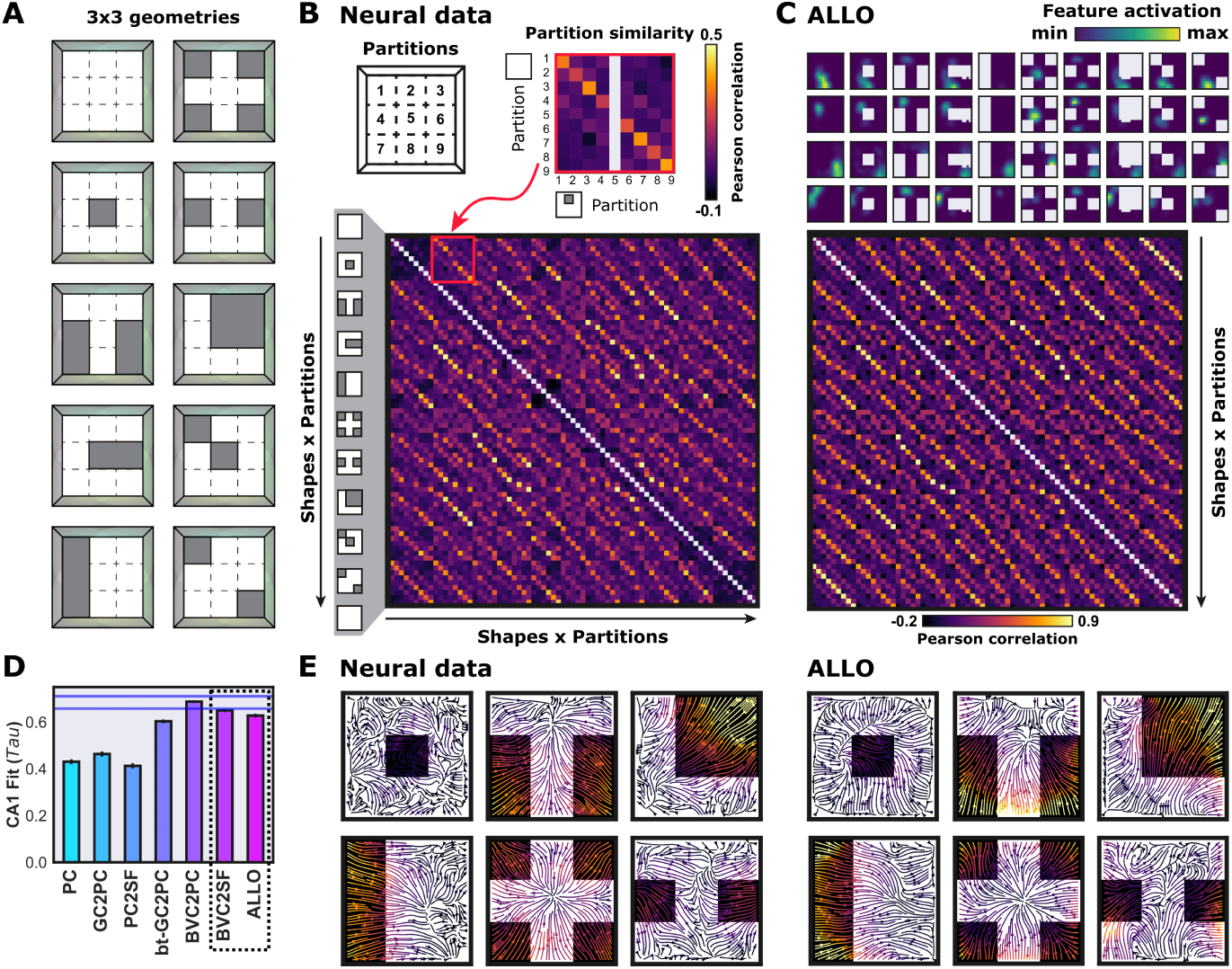
Remapping in emergent place-like representations is aligned to representations observed in CA1. (**A**) Environments used to evaluate remapping. Each frame corresponds to a different environment and each of them is defined by a 3 × 3 imagined grid of partitions, with some of them blocked by walls. During training, the model is exposed to a single environment; during evaluation, the model is exposed to three repeated sequences of 10 environments, starting with the open arena. (**B**) Representational similarity matrix (RSM) for the recorded CA1 neural data. Each partition of an environment is represented by a vector of 25 average activations, obtained from a finer grid inside each partition. The Pearson correlation between a vector pair determines the similarity of their corresponding partition pair. The RSM contains all the similarities for all the possible pairs of geometries and partitions. (**C**) Average activation of units in the model’s third MLP layer (above) and the respective RSM (below). (**D**) Rank-order correlations for different models (*33*). The rightmost bar corresponds to the proposed model; horizontal lines indicate noise thresholds from individuals. The dotted box compares the BVC2SF and the proposed models. BVC2SF suggests a similar connectivity between boundary vector cells and place cells. (**E**) Population vector flow maps for recordings (left), and the proposed model (right). Arrows depict shifts in population activity: each vector links a bin in a finer 15 × 15 imagined grid in the open arena to its most similar bin in another environment, based on Pearson correlation of the population vector.

From these activations, we computed the representational similarity between each pair of partitions in the 3 × 3 grid across all geometry pairs (Fig. 3C). To do so, we subdivided each tile into a 5 × 5 subgrid of bins and averaged the recorded activations across bins to produce a single vector per tile. We then measured similarity as the Pearson correlation between tile vectors, averaged across neurons (see fig. S8). This yielded one representational similarity matrix (RSM) per training run; these were averaged across runs to produce a single RSM summarizing the model’s representation.

To evaluate the model, we compared this average RSM to one derived from experimentally recorded CA1 population activity (*33*) (Fig. 3B). We flattened both matrices and computed their rank-order correlation (Fig. 3D). Notably, unlike the models in the original comparison, ours did not use the actual rodent trajectories for training but instead an approximation generated in a virtual 3D environment. Additionally, the other models were provided with allocentric spatial inputs, which our model lacked, placing it at a potential disadvantage. Despite this, our model achieved a rank order correlation of 0.628 ± 0.008, only 4.52% below the noise margin calculated from individual variability, and just 3.32% below the BVC2SF model (*24*), which, again, incorporates allocentric cues like distance to walls. A linear fit between RSM entries yielded an *R*^2^ of 0.827 ± 0.007 for our model, and 0.652 ± 0.010 for the BVC2SF model.

While high rank-order and linear correlations are encouraging, they may in part reflect a trivial preservation of activity within the allocentrically matching spatial bins across environments, as suggested by strong diagonal structure in the RSMs. To test whether our model captures meaningful remapping dynamics beyond this, we calculated rank order correlations for sub-matrices of the RSM that considered only different geometries or different partitions, obtaining similar values as alternative models (see fig. S9), and we generated a remapping stream plot following the methods used in (*33*). To obtain the stream plot, we subdivided the original open arena into a finer 15×15 grid, and represented each bin by the neural population vector that results from averaging all activations across neurons. For each environment, we used Pearson correlation to identify the most similar bin in all other environments, and used these bin pairings to construct a vector field depicting how representations shift with environmental changes (Fig. 3E). Strikingly, the resulting stream plots revealed consistent remapping patterns across geometries, including common systematic shifts in activity from blocked regions toward adjacent, unblocked areas.

### Model generality

The characteristic firing patterns of place and grid cells have been observed in the brains of various mammals (*51–53*) across multiple sensory modalities, including vision, olfaction, audition, and touch (*54, 55*). To evaluate the generality of spatial cell-type emergence in our eigenvector model, we trained multiple instances of the model on diverse observation modalities and systematically varied key hyperparameters of both the neural network architecture and the training algorithm.

We first evaluated the model’s generality across input modalities by training separate instances using four types of observations: one-hot encodings, 3D coordinates, and allocentric and egocentric pixel-based images (Fig. 4A). For the non-image inputs (one-hot and 3D coordinates), the convolutional module was replaced with a three-layer multilayer perceptron (MLP). After training, we assessed representational similarity using canonical correlation analysis (CCA) matrices instead of RSMs (Fig. 4B). This choice allowed us to account for potential mismatches introduced by eigenvalue multiplicity, using additional linear transformations to align the eigenvector approximations. Across input modalities, the resulting CCA matrices yielded Pearson correlations in the range [0.302, 0.652], indicating that the model can consistently reproduce parahippocampal–subicular representations across sensory domains. As a baseline comparison, we trained a variational autoencoder (VAE) (*56*) on the same data and computed CCA matrices on its latent representations (Fig. 4B), which produced substantially lower Pearson correlations in the range [0.007, 0.158].

**Figure 4:**
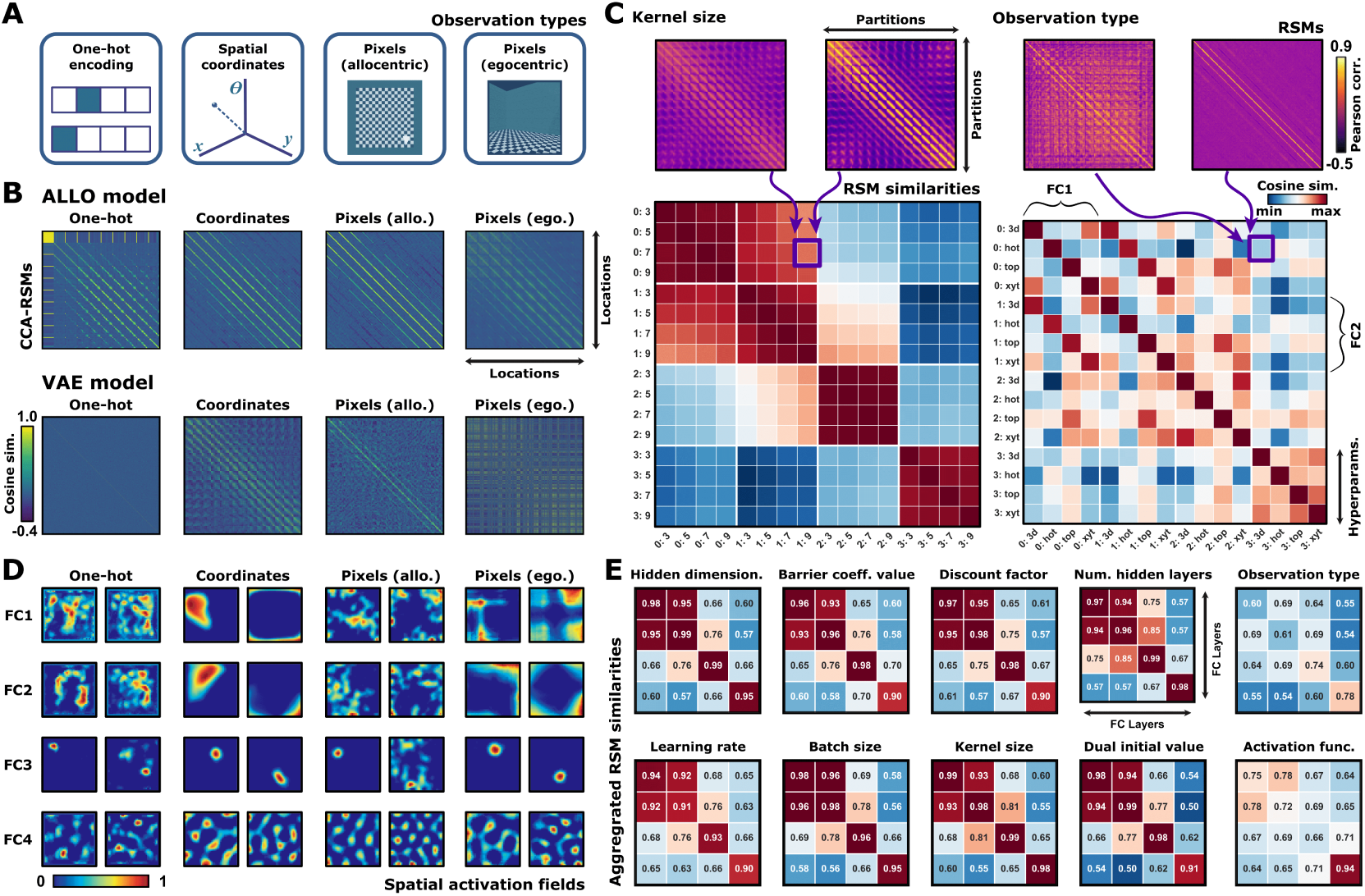
Emergent representations mirroring the hippocampal-entorhinal system are observation dependent but robust across hyperparameter tunings. (**A**) Observation types used as inputs: one-hot encodings representing a discrete 3D grid over (*x*, *y*, θ) (location and head direction), continuous [*x*, *y*, θ] vectors, pixel-based allocentric (top-down) views, and pixel-based egocentric views. (**B**) Canonical correlation analysis (CCA) matrices comparing model output features for each observation type. The matrices correspond to the proposed model (above) and an alternative VAE model trained on the same inputs (below). Each matrix compares the representation of a single observation type with itself and each entry contains the cosine similarity for a pair of locations after the CCA transformation. Only 25% of the locations is shown in the case of the proposed model for ease of visualization. (**C**) Cosine similarity matrices across hyperparameter values for two settings: kernel size in the convolutional module (left) and observation type (right). An entry contains the cosine similarity between two representational similarity analysis (RSM) matrices, each corresponding to a hyperparameter value and a specific layer. Each entry in an RSM reflects similarity between population vectors in corresponding tiles of a 15 × 15 partition of the open arena. (**D**) Two example spatial activation maps across layers for each observation type. (**E**) Average cosine similarity matrices across all values for each hyperparameter. Each entry compares the representational similarity between layers.

Finally, we evaluated the representational similarity between layers across a range of hyperparameter values (see (*32*) for details). For ten different hyperparameters, we trained multiple model instances (ten random seeds per value, per hyperparameter) and computed an RSM for each value-layer pair (Fig. 4C, above). In particular, the hyperparameters include one dataset (observation type), four architectural (e.g., activation function), and five algorithmic (e.g., learning rate) choices (Fig. 4E). We then calculated the cosine similarity between these RSMs and aggregated them by layer, resulting in a cosine similarity matrix for each hyperparameter (Fig. 4C, below). To aid visualization, we further compressed each matrix by averaging over all layer pairs (Fig. 4E; see fig. S10 for remaining full matrices).

With the exception of observation type and activation function, all hyperparameters produced highly diagonal similarity matrices, indicating that each layer maintained distinct and consistent representations, suggesting that the units’ representations, across a wide range of model configurations, consistently mirror those hierarchically observed in the hippocampal-entorhinal system, as shown in Figure 2. In the case of 3D coordinate inputs, place cells did emerge, and early-layer activations covered broad spatial regions with some border preference. However, these patterns were qualitatively different from those typically found in the subiculum. For one-hot encoded and top-down image observations, units resembling place cells often exhibited multiple peaks (typically between one and four) rather than a single localized field. Likewise, units resembling subicular cells displayed irregular, dispersed activity with no clear preference for borders or corners (Fig. 4D).

Taken together, these results demonstrate the robustness of the proposed method across a wide range of hyperparameters, including variations in the network architecture itself. Notably, although specialization across layers was observed for various observation types, hippocampal- and subiculum-like activity only closely matched biological data when the model was trained with biologically plausible set of observations (optical flow). This finding shows that our model can solve the localization problem using only constrained, partially observable egocentric inputs without access to explicit allocentric information. It also suggests that the neural representations observed in the brain may similarly reflect a structured solution composed of multiple specialized cell types. This echoes findings in human object recognition: the richness of visual input is predictive of neural alignment in the occipitotemporal cortex (*57*), and head-mounted eye-tracking studies in toddlers show that training contrastive models on limited visual fields improves object recognition performance (*58*). In contrast, when allocentric information is available, many representational shortcuts may likely enable localization, potentially bypassing the need for the compositional structure observed in the hippocampal-entorhinal system.

## Discussion

The computational roles and inter-dependencies of spatially tuned neurons in the hippocampal–entorhinal system remain a subject of debate, with contrasting predictions of several models and interpretations of empirical findings. Some studies have argued that grid cells are necessary for the emergence (*2,26,59*) and stability (*27*) of place cells, while others propose that spatial representations of place cells are construed from combinations of boundary vector cells (*14,15,21,24*). Conversely, some models posit that grid cells require input from place and head-direction cells (*3,6,60*), or are formed from combinations of band cells (*35*). Our results offer a unifying framework in which spatially tuned cells in the hippocampal–entorhinal system emerge hierarchically and in parallel, relating sensory observations to navigational demands through predictive learning. Criti-cally, the model suggests a bidirectional dependence: earlier layers construct increasingly abstract representations that support higher-level predictive coding, while error signals from deeper layers constrain and refine the earlier representations, in agreement with empirical observations (*61*). This predicted interdependence and sequential organization of representations in our model can be leveraged to generate new predictions on the causal relationship among representations in the entorhinal-hippocampal system.

Our results also suggest that diverse cell types in the parahippocampal–subicular region, like place, band, grid, border, and corner cells, may serve a common computational purpose: approximating the spectral structure of environmental dynamics. Extending earlier proposals (*19, 35, 59*), we show that these cells encode eigenvectors of the successor representation (SR) matrix (*19*) (the SR operator). This spectral view also clarifies the distinct roles of cells in different subregions: while subicular boundary vector and corner cells (*46*) express activity patterns similar to border and center-distance cells in the medial entorhinal cortex, their computational role is distinct: subicular cells provide information necessary to build a map from egocentric inputs, while medial entorhinal cells express the spatial coordinates of this map. This interpretation helps to reconcile seemingly conflicting interpretations of experimental results. For example, although place cells depend on input from the medial entorhinal cortex (*26, 59*), they can still emerge after grid cells are disrupted through septal inactivation (*62*). Since grid cells but not other parahippocampal inputs are affected by this precedure, CA1 could continue to receive sufficient spectral input for place cell emergence. Similarly, since other regions such as the pre- and parasubiculum contain cells with grid-like responses and project strongly to CA1 (*63*), our model predicts that place cells would still emerge with inputs from these regions, and could explain why large subpopulations of hippocampal cells remain spatially selective despite complete removal of the medial entorhinal cortex (*27*).

The emergence of functional cell types in the hippocampal–entorhinal system has previously been demonstrated in models based on path integration leveraging deep recurrent architectures (*3, 5, 6, 23, 64*). In such models, grid cells arise from error correction in position estimation (or a latent version (*5*)), and place and head-direction cells are the first to emerge to support the recurrent mechanisms in the tuning of speed cells (*3, 6, 23*). However, these approaches often rely on bespoke training frameworks, do not explain the emergence of pre-existing cells, and do not align with the experimentally observed dependence of place cells on grid cells (*26*). By contrast, our eigenvector model agrees with a body of work suggesting that path integration and attractor dynamics are not required for the emergence of grid-like representations (*12, 65*). Instead, our results align with the view that spectral structure in sensory input is sufficient to induce neurobiologically plausible spatial cell types (*12, 65–67*). Moreover, while prior work has shown that grid cell emergence in path integration models is highly sensitive to narrow ranges of hyperparameters (*6, 65*), we found that the activity mirroring the hierarchy in the hippocampal-entorhinal system emerged robustly in our model across a wide range of settings.

Our model exhibits several properties that contribute to its biological plausibility. First, the only consistent dependencies we observed were on the type of nonlinearity, where a non-negative activation function (ReLU) and on the use of egocentric observations were crucial. Both constraints are neurobiologically grounded: non-negativity aligns with the firing properties of real neurons (*68*), and egocentric input reflects the partial and embodied nature of an animal’s sensory experience. Beyond these architectural features, the spectral graph learning algorithm we employ (*30*) also depends on two memory-related mechanisms: a buffer to store past observations and a replay process that samples from this buffer with probabilities shaped by the successor representation (SR). These requirements are compatible with what is known about the hippocampus, a brain region central to episodic memory (*27, 69*). Findings suggest that the population activity of place cells exhibits transient patterns that function as “barcodes” for specific experiences, which are replayed during memory-related tasks (*70–73*). Moreover, a recurrent model accounting for barcode formation has been shown to encode SR-like probabilities in its synaptic weights (*74*), providing a plausible mechanism through which learned transition statistics guide replay. Taken together, these elements suggest that our model’s learning dynamics are not only computationally effective but also conceptually aligned with recent experimental observations in the hippocampal-entorhinal system.

The results we presented also add to a growing body of evidence positioning SR- and Laplacian-based methods as a promising direction in computational reinforcement learning. The core ideas behind ALLO have previously been used to construct temporal abstractions, enabling effective exploration and control in high-dimensional tasks (*75*). More broadly, spectral and contrastive representations have been leveraged for skill discovery (*76,77*), exploration (*78*), planning in abstract metric spaces (*79, 80*), and zero-shot generalization (*81–85*). These advances mirror behavioural evidence from human studies, where the SR has been proposed as a computational substrate for semi-flexible choice (*86*) and for modelling human decision-making across multiple tasks (*87, 88*). Taken together, this body of work suggests that the SR (and its eigenspectrum) may serve as a natural foundation not only for spatial and temporal abstraction (*89*), but for sequential decision-making more broadly.

Beyond the implications discussed for both neuroscience and computational reinforcement learning, our model and methodology open new avenues for research at the intersection of artificial intelligence and neuroscience. By training on egocentric, biologically plausible sensory observations, we can now simulate diverse functional properties observed in the entorhinal–hippocampal system, enabling the generation of rich sets of hypotheses in silico, which can stimulate new experimental evaluation in vivo. Importantly, because the eigenvector model is input-agnostic, our model can be extended to any sensory modality and task space, such as abstract symbols or sound, raising the possibility of explaining—or even predicting—diverse emergence of other non-spatial cell types recently observed in the hippocampal-entorhinal system (*90–92*).

## Funding

D. G., M. B., and M. C. M. were funded in part by the Natural Sciences and Engineering Research Council of Canada (NSERC) and the Canada CIFAR AI Chair Program. M. B. was funded in part by the Google DeepMind Chair in AI. This research was enabled in part by the computational resources provided by the University of Alberta, the Digital Research Alliance of Canada.

## Author contributions

Conceptualization, D. G. and M. C. M.; methodology, D. G., J. Q. L., and M. C. M.; investigation, D. G.; visualization, D. G., M. B., J. Q. L., and M. C. M.; funding acquisition, M. B. and M. C. M.; project administration, M. C. M.; supervision, M. B., J.Q.L, and M.C.M; writing– original draft, D. G. and M. C. M.; writing– review & editing, D. G., M. B., J. Q. L., and M. C. M.

## Competing interests

There are no competing interests to declare.

## Data and materials availability

Data and code to reproduce figures will be publicly available at the time of publication.

## Supplementary Materials for

### Materials and Methods

#### Experimental setup

##### Environments

To simulate rodent trajectories, we made use of the MiniWorld library (*93*), which supports the definition and visualization of 3D environments with pixel-based observations. In particular, it includes both allocentric (top-down view) and egocentric perspectives.

These environments are standard reinforcement learning environments with discrete time steps. Hence, we simulated the movement of the rat by having a stochastic policy that selects one action from a set of 6 possible actions after receiving each observation: move in a straight line, move backwards, rotate the head to the left or to the right, and rotate to the left or right while simultaneously moving along the head-direction. The distance and orientation displacements resulting from these actions were sampled uniformly from ranges chosen to generate realistic trajectories. Also, the displacements were further perturbed by adding rotational uniform noise to translational ones and vice versa.

##### Exploratory policy

The stochastic policy used to select actions is composed of three subpolicies that are executed in different cases: when a wall is close and perpendicular to the agent, when it is close, but parallel, and when there are no nearby walls. In the first case, the policy avoids the wall by rotating the head towards the direction that makes the agent most parallel to the wall. In the second case, with some probability, the policy chooses to move forward, mimicking the preference of rats to move near walls, and otherwise uniformly randomly. In the last case, the policy also chooses uniformly.

##### Visual cues

Without any memory mechanism, like recurrence, there is no way for a model to deduce the allocentric configuration from purely egocentric views. Inspired by animal experiments that typically signal orientation through cue cards on walls, we included four sources of light of different colours in the square open arena (see Fig. 1A,B). Each of them points from the middle of one wall to its parallel wall. The same four sources were included in all of the environments considered for remapping (see Fig. 3A) and for the different topologies, as we found them sufficient for the emergence of the expected periodic patterns.

#### Learning algorithm

##### The Laplacian operator

We refer to the configuration of an agent and its environment, the state, as a vector *s* in a vector space S. The agent interacts with the environment by executing actions, chosen from a set *A*, by means of a stochastic policy π : *S* → Δ(*A*), where Δ(*A*) is the simplex over the set *A*. The environment evolves following the transition map *M* : *S* × *A* → Δ(*S*) that specifies how likely it is to visit a future state after a state-action pair. This interaction determines a probability law *ℙ_π,M_* from which random variables and expectations of them can be defined in a sound manner. In particular, *P*_π_ : *ℝ^S^* → *ℝ^S^* is the transition dynamics operator defined as the expected value of the transition map: 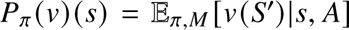, where *v*; ∈ *ℝ^S^* is a function over *S*. In the finite-dimensional case, this is the same as saying that *P*_π_ is a matrix that results from the tensor product *M* × π and encodes the probabilities of transitioning from one state *s* to a future state *s*^′^, considering that the agent follows π. Hence, the product *P*_π_*v*; is equivalent to the expected value of *v*; after one transition. The SR operator is defined as the geometric series of the discounted dynamics operator γ*P*_π_, i.e., 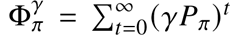. The SR captures the expected occupancy of a future state *s*^′^ starting from another state *s*, but considering all the possible future transitions, and not only the next one. The weighting of the future occupancies is discounted by the discount factor γ. Moreover, the Laplacian operator is typically defined as the difference 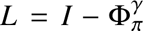 (typically with γ = 0, which implies *L* = *I* − *P*_π_), where *I* : *ℝ^S^* → *ℝ^S^* is the identity operator. These operator connects the reinforcement learning framework with graph theory, where the Laplacian operator is typically used to evaluate functions over graphs (in this case, the state graph specified implicitly by the dynamics) and to determine graph properties related to connectivity, bottlenecks, and traversal times.

##### Augmented Lagrangian Laplacian Objective (ALLO)

Finding the eigendecomposition of the Laplacian operator has different important uses in computational reinforcement learning, like exploration and skill discovery, and in graph theory, like graph visualization. In our case, the interest lies in the eigenvectors exhibiting similar periodic patterns as those observed in grid cells. Performing this eigendecomposition in the more general abstract setting, where *S* is possibly uncountable, is unfeasible via standard linear algebra solutions. Instead, it can be shown (*30*) that the eigenvectors of the Laplacian are a minimizer of the following constrained optimization problem:

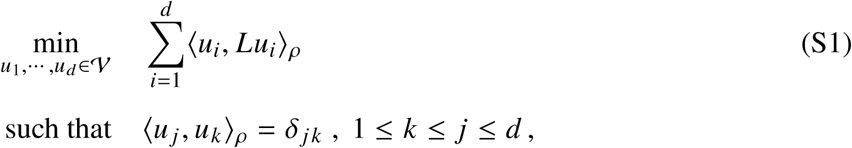

where *u*_*i*_ ∈ V ⊂ *ℝ^S^* is a function over 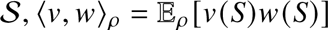 is the inner product in *S* induced by the distribution ρ, and δ _*j k*_ is the Kronecker delta, which is equal to 1 when *j* = *k* and otherwise 0. The constraints ensure that the solution functions are orthonormal, and the inner product in the cost function selects for vectors that minimize the projection of themselves with their transformation after the application of the Laplacian. Since the eigenvectors form a basis, the transformation consists of a combination of eigenvectors scaled by the eigenvalues, and then it is intuitive that the projection is minimized by the eigenvectors with lower eigenvalues. Specifically, the solution corresponds to the *d* eigenvectors of the Laplacian *L* with associated smallest eigenvalues, which is equivalent to the eigenvectors of the SR 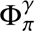 with largest eigenvalues.

In practice, however, finding the solution of the optimization problem requires gradient descent based solutions whose dynamics may not necessarily converge to the desired minimizer. The Augmented Lagrangian Laplacian Objective (ALLO) (*30*) is an algorithm that ensures convergence to the desired eigensystem, by the simultaneous iterative maximization and minimization of the objective of the same name:

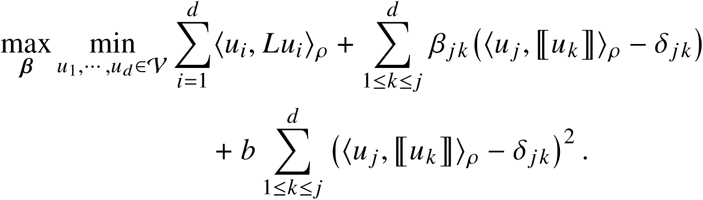

Here, ⟦·⟧ refers to the stop-gradient operator, which indicates that, when following gradient descent dynamics, the real gradient of the objective should not be used. Instead, when calculating derivatives, whatever is inside the operator is assumed to be constant. The introduction of the linear constraints, (⟨*u* _*j*_, ⟦*u*_*k*_ ⟧⟩_ρ_ −δ_*j k*_), in the cost function, multiplied by the dual parameters, ***β*** = [β_*j k*_] _*j k*_, ensures that the eigenvectors are an equilibrium point of the resulting dynamics, and the squared constraints, (⟨*u* _*j*_, ⟦*u*_*k*_ ⟧⟩_ρ_ − δ _*j k*_)^2^, multiplied by an appropriate barrier coefficient *b* make the eigenvectors, ordered by their eigenvalues, the only stable equilibrium point.

##### SR-based Monte Carlo sampling

In realistic problems where observations correspond to optical flow, the previous max-min objective cannot be computed explicitly. The Laplacian operator encoding the transition dynamics is unknown and, even if it was known, it has a large dimensionality. The deep learning approach consists of generating Monte Carlo approximations of the partial gradients with respect to the primal (the state features *u*_*i*_) and dual parameters and updating these parameters with the approximations. In this particular case, it can be proven (*30*) that the objective can be rewritten as:

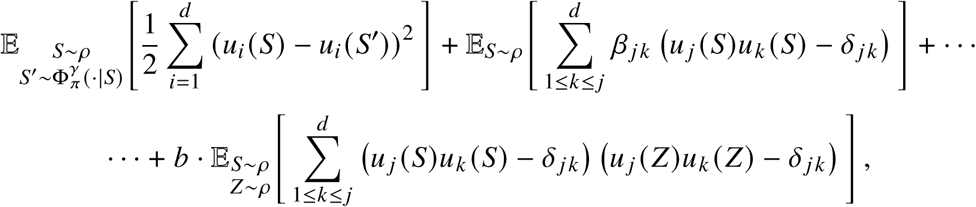

which consists of pure expectations that can be approximated by sampling uncorrelated states *s*, *z* from some chosen distribution ρ and future states from the environment’s SR operator 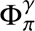.

To replicate this sampling procedure in practice, we use an episodic replay buffer—essentially, an array, which we fill with episodic trajectories of observations. At each optimization step, a batch of triples of states 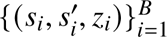 of size *B* is generated. We sample *s*_*i*_ and *z*_*i*_ uniformly *i i*=1 from episodes and steps, and, for each *s*_*i*_, we sample *s*’ = *s_i=+k_* by sampling a future index *k* that follows a geometric distribution with parameter γ. This is equivalent to sampling states from the stationary state distribution induced by the exploratory policy interacting with the environment (or a truncated version, because of a maximum imposed length of steps) and future states from the effective discounted state distribution determined by the SR operator 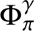.

##### Episodic training

The model training consists of the collection of 100 trajectories of length 10, 000 and the simultaneous use of this data to update the model parameters according to ALLO. Trajectories are generated simulating the interaction of the exploratory policy with the MiniWorld environments. After an initial collection of episodes (5%), the model is updated for 200,000 steps. The rest of the trajectories is generated in a uniformly spaced manner throughout updates (1 trajectory every ∼ 21, 000 steps). These numbers were chosen for computational reasons. In earlier experiments, we collected 100% of data before training and noticed no significant changes.

##### Deep learning model

The model we use is a deep neural network *u*_θ_ : O → *ℝ*^*d*^ that maps observations in the space O (R^60×80×3^ in the case of egocentric views) to a feature space of dimensionality *d*. In this way, the *i*−th dimension *u*_θ,*i*_ (*o*) evaluated at an observation *o* ∈ O corresponds to the approximation of *u*_*i*_ (*s*) in the ALLO objective (this assumes that states correspond 1-1 to observations, explaining the need for the visual cues). Hence, after training, *u*_θ,*i*_ encodes the *i*−th eigenvector of the Laplacian operator associated to the SR. In addition, we use a lower triangular matrix ***β*** ∈ *ℝ*^*d*×*d*^ to store the dual parameters.

The network is composed of a convolutional module Conv_θC_ followed by an MLP module MLP_θMLP_ : *u*_θ_ = MLP_θMLP_ ◦ Conv_θC_. The convolutional module has 3 layers with 16 channels, a stride of 2 for the first two and 1 for the last, no pooling, same padding, and a kernel size of 3, except for the experiment where the kernel size is varied.

To train the model *u*_θ_, we use the Adam optimizer with learning rate 3 × 10^−4^. For the duals, we use vanilla stochastic gradient ascent instead of Adam, with the same learning rate, since we observed that the latter made the dual dynamics slower than needed.

#### Cell classification

To classify neurons in the deep neural network, we place the agent in each (*x*, *y*) location and head orientation θ in a uniform 3D grid of size 40 × 40 × 12 and register the output of all neurons in the MLP module. The resultant activations are averaged across orientations and locations, obtaining an activation field of size 40 × 40 and a direction-tuning signal of length 12 for each neuron. Then, we assess different scores defined in the literature for both fields and direction signals. These scores are compared against some fixed threshold or a percentile obtained from spatially shuffled versions of the activations.

##### Spatial shuffling

Following (*35*), we generate random shuffles of the Voronoi tesselations of the activation fields. For each neuron, we estimate the local peaks in its activation field and then assign each pixel in the field to the closest peak, generating a tesselation made out of polygons. These polygons are randomly relocated in space and rotated with respect to their centroids. For pixels in the intersection of polygons and outside of boundaries, due to transformations, we individually and randomly place them in regions that the transformed polygons do not cover. In this manner, the norm of the field is not modified.

##### Grid cells

To identify grid cells, we follow the methodology outlined in (*34*). Specifically, we compute the autocorrelograms of the activation fields and identify all the peaks in them, defined as the locations where the value is maximal across a neighbourhood of 1 pixel radius and is above a threshold of 0.2 times the maximum autocorrelation. From the peaks, we extract a set of annuli, centred at the origin, whose external and internal radii are chosen to include 6 peaks and to exclude the central peak, respectively. In case that there are fewer than 6 non-central peaks, we include all of them. The annuli are rotated across multiple angles, including 30°, 60°, 90°, 120°, and 150°, and the Pearson correlation is computed with respect to the original annuli. An activation field that has perfect hexagonal symmetry will have 6 six peaks separated by 60°. Hence, the correlation across rotations would exhibit a perfect sinusoid with a period of 60°, with a maximum of 1 at 0°and 60°and a minimum of 0 at 30°. Correspondingly, the gridness score is defined as the minimum gap between the maximum and minimum correlations. For the maximum, we consider the minimum of the correlations between 60°and 120°, and for the minimum, the maximum at 30°, 90°, and 150°. We classify a neuron as a grid cell if its gridness score is above 0.0.

##### Head-direction cells

For each neuron *i*, we compute the magnitude of its average Rayleigh vector 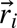, where the Rayleigh vector at angle θ is defined as 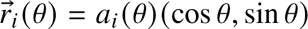, with *a*_*i*_ (θ) being the maximum-normalized directional-tuning of neuron *i* at θ. Hence, 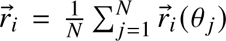, with *N* = 12 and each angle corresponding to equally spaced directions from 0°to 360°. We define the directionality score as the difference between the magnitude 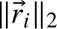 and the 95th percentile of similar magnitudes for angular permutations of the original directional-tuning signals, similar to the approach taken in (*35*). The permutations were generated uniformly, one by one, until the 95th percentile of the magnitude only changed by a small amount. A neuron is classified as a head-direction cell if its directionality score is above 0.0.

##### Periodic cells

To determine if a neuron exhibits periodicity, we generate spatial shufflings of its activation field and compute their Fourier transforms until the 95th percentile of the maximum Fourier component varies less than a small threshold. We then compute the Fourier transform of the original activation field. A neuron is classified as a periodic cell if the magnitude of its Fourier spectrum has a peak above the 95th percentile of the shuffled spectra, in line with previous work (*35*).

##### Border and boundary vector cells

We calculate the borderscore of a neuron to assess its preference for borders, as defined in earlier works (*40, 46*). First, we identify all pixels in the activation field that are above a threshold of 0.3 times the maximum activation. Then, we compute two quantities: the maximum fraction of pixels along any border that belong to the same connected neighbourhood (CM); and the mean weighted normalized distance of the high-activation pixels to the closest boundary (DM), where the normalized activations of those same pixels are used for the weighting and the distance is normalized with respect to the centroid of the environment. The borderscore is defined as the ratio CM − DM/CM + DM and a neuron is classified as a border or boundary vector cell if its borderscore is above 0.45. Intuitively, this metric is the gap between the boundary and the central activation, with a value of 1 for an activation that only happens in the border, and -1 for one that only happens in the centroid. The only distinction we make between border and boundary vector cells is that we use the former to refer to neurons in the output layer of the model, while the latter refers to hidden layers. This choice mirrors the use of the metric to identify both cell types, depending on whether they are located in the medial entorhinal cortex (*40*) or the subiculum (*46*).

##### Corner cells

To identify corner cells, we use the cornerscore introduced in (*46*). Similarly to the borderscore calculation, we identify connected neighbourhoods of high-activation pixels and then compute two opposite quantities that capture distances to the centroid of the environment and its corners. Specifically, the cornerscore associated to a neighbourhood is defined as the ratio *d*_1_ − *d*_2_/*d*_1_ + *d*2, where *d*_1_ is the distance between the pixel with maximal activation in the neighbourhood and the centroid, and *d*_2_ is the distance between this same pixel and the closest corner. The cornerscore is defined as the average of the neighbourhood cornerscores minus a penalization term. Denoting *k* as the number of corners, the average is calculated over the *k* neighbourhoods with the largest cornerscore, and the penalization for each additional neighbourhood *i* is (1 − cornerscore_*i*_)/*k*. The cornerscore is maximal with 1 when all activation happens in the corners, and minimal with −1, when the only activation happens at the centroid. We use the same threshold of 0.3 to identify high-value pixels. A neuron is classified as a corner cell when the cornerscore is above 0.5.

#### Remapping

##### CA1 Neural dat

We use publicly available data (https://zenodo.org/records/14867736) consisting of a dictionary that registers the recorded locations and simultaneous neural firing rates of hundreds of CA1 neurons in 4 male and 3 female rats. The recordings correspond to 31 sessions of 40 minutes, where each session was carried out on a different day in a different geometry. The geometries were chosen following a randomized repeating sequence of the 10 possible geometries, starting and ending with the open arena.

##### Training

We train a separate instance of the deep learning model for each of the 10 geometries, as specified previously. In particular, this means that we use the exploratory policy to generate trajectories, and we do not make use of the recorded locations. Also, the 10 instances are trained independently, without considering any of the sequences of environments that the rats experienced. *Model data:* To benchmark our model against the existing CA1 data, we simulated the trajectories described by the recorded locations across the 31 sessions for one of the seven rats. Since only (*x*, *y*) coordinates are available, we inferred the head orientation from the displacement vectors between two successive positions. Then, we recorded the activations of the third fully connected layer of each independent instance of the model, in the same format as the real recorded CA1 firing rates.

##### Data analysis

We use the code released by Lee et al. (*33*) to create spatial activation maps for each day as well as for their analysis, including the calculation of the representational similarity matrices (RSMs), the rank order correlation between pairs of RSMs, and the population vector flow maps.

We also generated spatial activation maps by placing the agent in each possible location of a grid in the (*x*, *y*, θ) space for each possible model-geometry pair and directly calculated the corresponding RSM. That is, we calculated an RSM without using the recorded trajectories or sequences of environments. Despite this, we obtained a similar rank-order correlation of 0.59 between the model and the CA1 data.

#### VAE model

We consider the β−VAE (variational autoencoder) algorithm to train an encoder-decoder deep learning architecture. The encoder network *f*_θ_ : *O* → *ℝ*^2*d*^ maps observations to latent representation parameters, and the decoder network *g*_θ_ : *ℝ*^*d*^ → O reconstructs observations from sampled latent codes.

##### Encoder and decoder architectures

The encoder architecture is the same as the one used for the deep learning model that learns the Laplacian eigenvectors. The only difference is that the final layer generates a vector of size 2*d* = 80 with concatenated mean and log-variance parameters [μ, log σ] ∈ *ℝ*^2*d*^. The latent representation corresponds to a normal distribution, and samples are obtained by adding the mean to a randomly generated Gaussian noise ε that is scaled by the standard deviation.

The decoder mirrors the encoder architecture, in reverse. For visual inputs, it uses an MLP to map from the *d*-dimensional latent space to an intermediate representation, which is then reshaped and processed through transpose convolution layers, implemented as nearest-neighbour upsampling followed by convolution to avoid checkerboard artifacts. For non-visual inputs, the network consists just of an MLP. The decoder’s final activation depends on the observation type: ReLU for images, tanh for coordinates, and identity for one-hot observations.

##### Loss function

The training objective combines a reconstruction loss and a KL divergence regularization, weighted by the hyperparameter β:

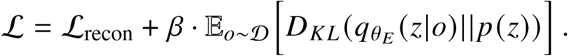

The reconstruction loss varies by observation type to accommodate different data characteristics.

For coordinates and visual observations, we use the Huber loss with parameter 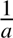

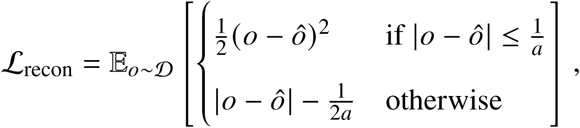

where 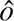 represents the reconstruction for observation *o*.

For one-hot observations, we employ a balanced binary cross-entropy loss that selectively focuses on the active entry and one randomly selected inactive entry to address class imbalance:

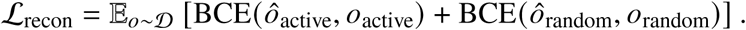

In addition, the KL divergence term regularizes the latent space toward a standard Gaussian prior: 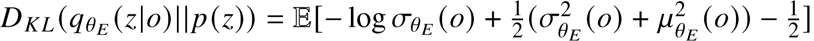

##### Background separation

The reconstruction of top-view images can be challenging since the largest fraction of an image corresponds to a static background *b*. Considering this, we add an estimation of the background to the decoder output 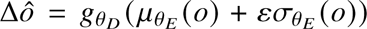. That is, the reconstructions are 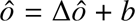, forcing the model to focus on learning to model variations, rather than full images.

We estimate the background by performing an adaptive averaging of all collected observations, representing the most frequent visual patterns in the environment. Specifically, we sweep over the existing dataset, one sample at a time, rejecting those pixels that are estimated to be outliers because they are too far from their corresponding current estimated mean.

**Figure S1:**
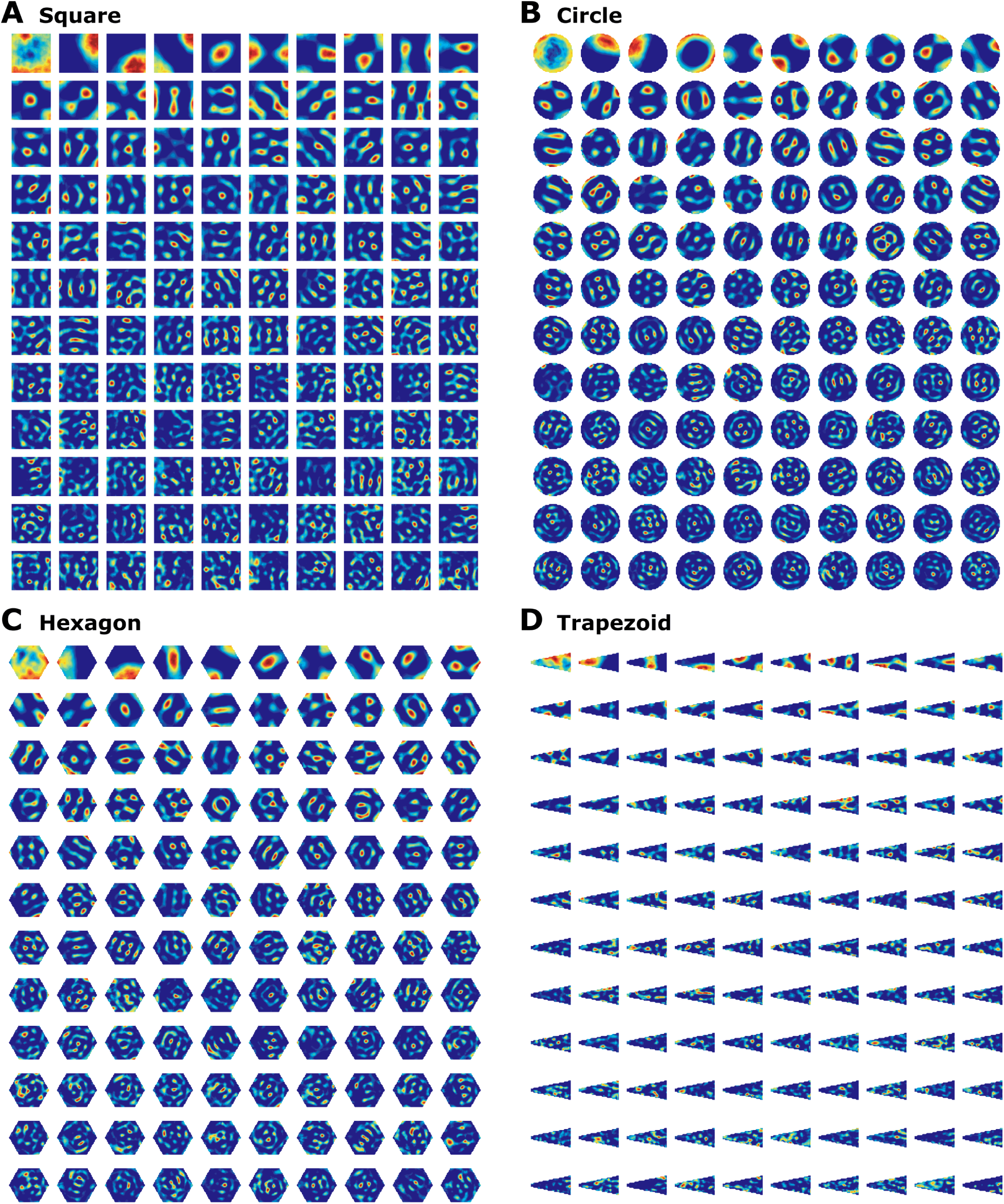
Representations learned in open arenas with different topologies. Clipped outputs of deep learning models trained with the ALLO objective for (**A**) a squared, (**B**) circular, (**C**) hexagonal, and (**D**) trapezoidal empty rooms.

**Figure S2:**
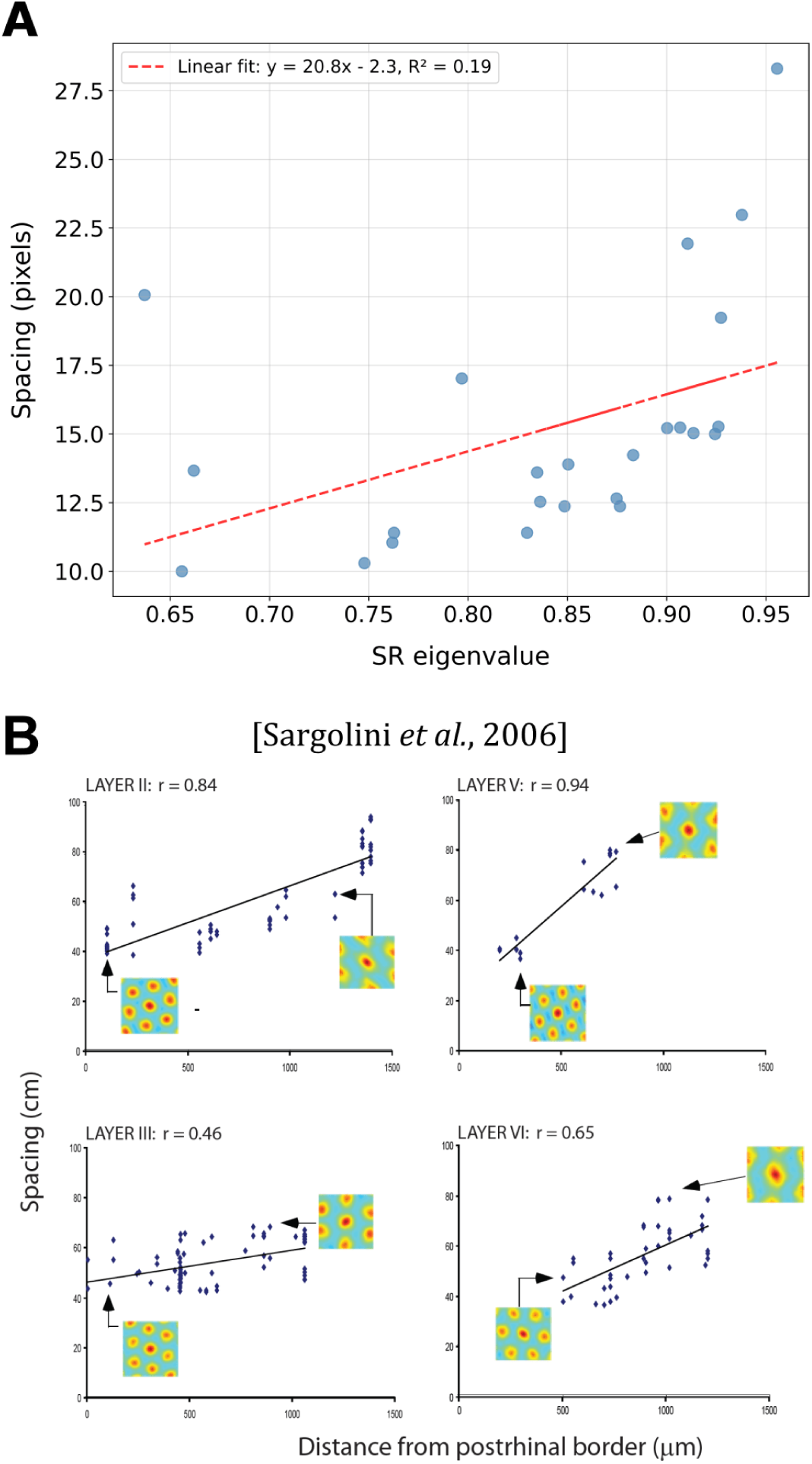
Spacing presents a monotonic behaviour in grid cells. Comparison between model and recorded neural grid spacing, defined as the mean distance between the central and the closest 6 peaks (or the maximum number if fewer than 6). (**A**) Grid spacing as a function of the corresponding eigenvalue in the output features of the model identified as grid cells. The eigenvalues are obtained as a byproduct of optimizing the ALLO objective and correspond to the dual variables β_*ii*_. (**B**) Grid spacing as a function of distance to the postrhinal border for different layers of the medial entorhinal cortex. Taken from the Supporting Online Material by Sargolini et al. (*34*).

**Figure S3:**
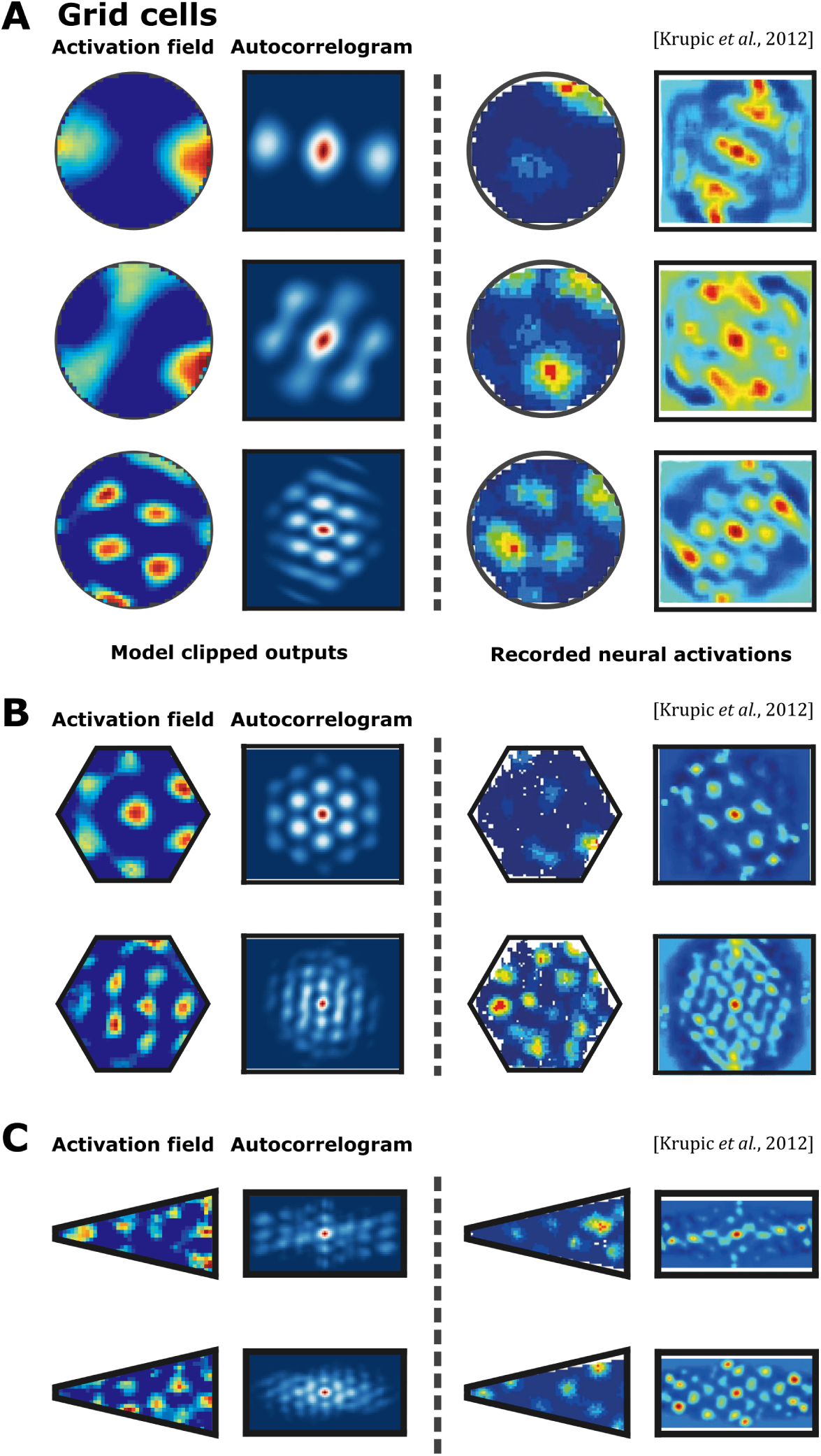
Grid cell density increases on different topologies. Comparison between model features (left columns) and firing patterns observed in rodent studies (right columns). For each model feature, the spatial activation map (second-left column) and autocorrelogram (second-left column) are shown. (**A**) Circular topology. (**B**) Hexagonal topology. (**C**) Trapezoidal topology.

**Figure S4:**
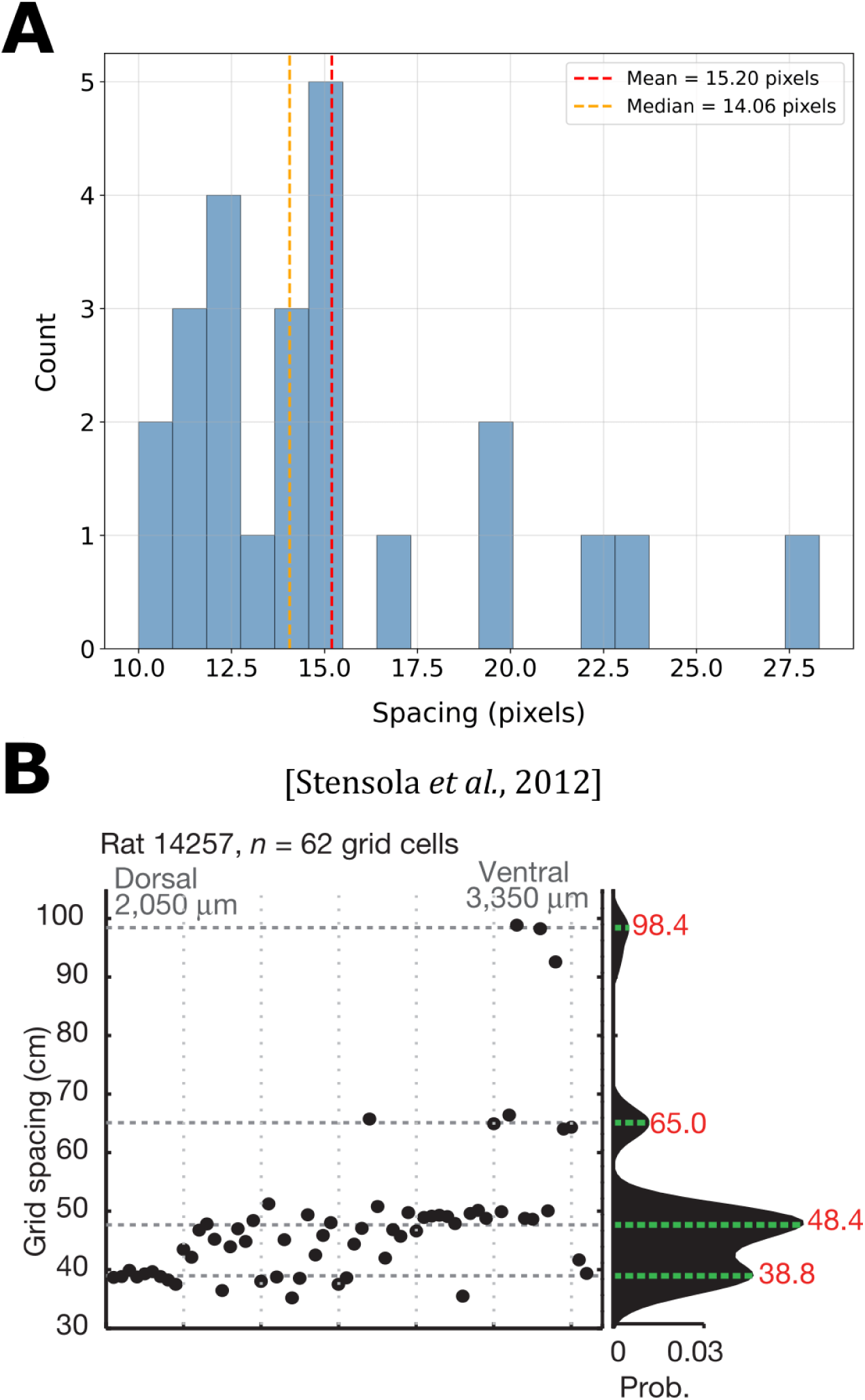
Grid spacing histogram. Comparison between model and recorded neural grid spacing distributions, where spacing is defined as the mean distance between the central and the closest 6 peaks (or the maximum number if fewer than 6). (**A**) Histogram of grid spacing for the output features of the model identified as grid cells. (**B**) Grid spacing density model displaying four characteristic peaks in the medial entorhinal cortex. Taken from (*39*).

**Figure S5:**
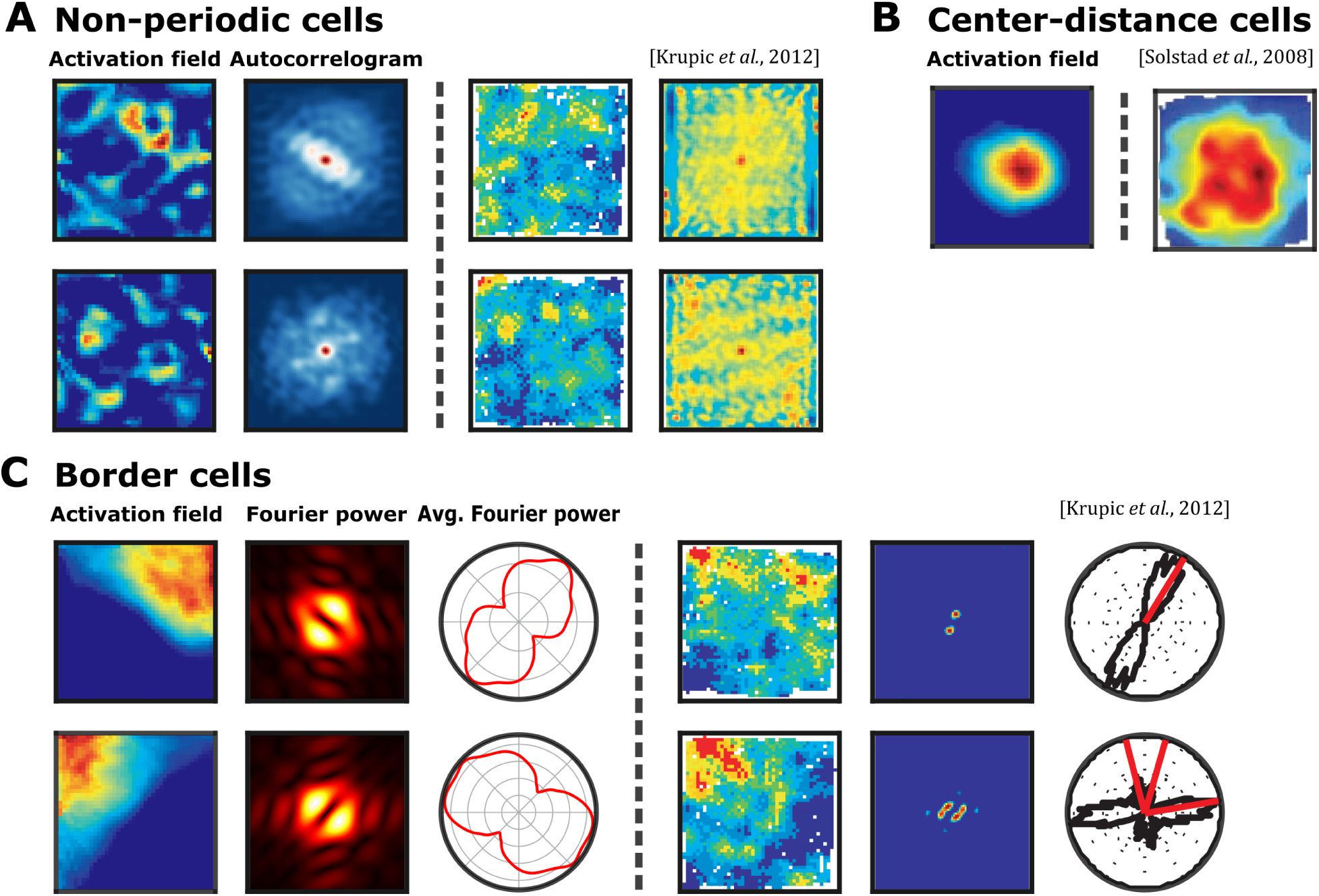
Additional entorhinal, pre- and parasubicular emergent cells. Comparison between model features (left columns) and firing patterns observed in rodent studies (right columns). For each model feature, the spatial activation map, autocorrelogram (second-left column), and polar plot are shown (third-left column) where applicable. (**A**) Non-periodic spatially tuned cells recorded in the entorhinal cortex. (**B**) Center-distance cells. (**C**) Border cells activate along walls and exhibit few Fourier components (the second column shows magnitudes of Fourier spectrograms, and the third column shows the radial averages of Fourier powers).

**Figure S6:**
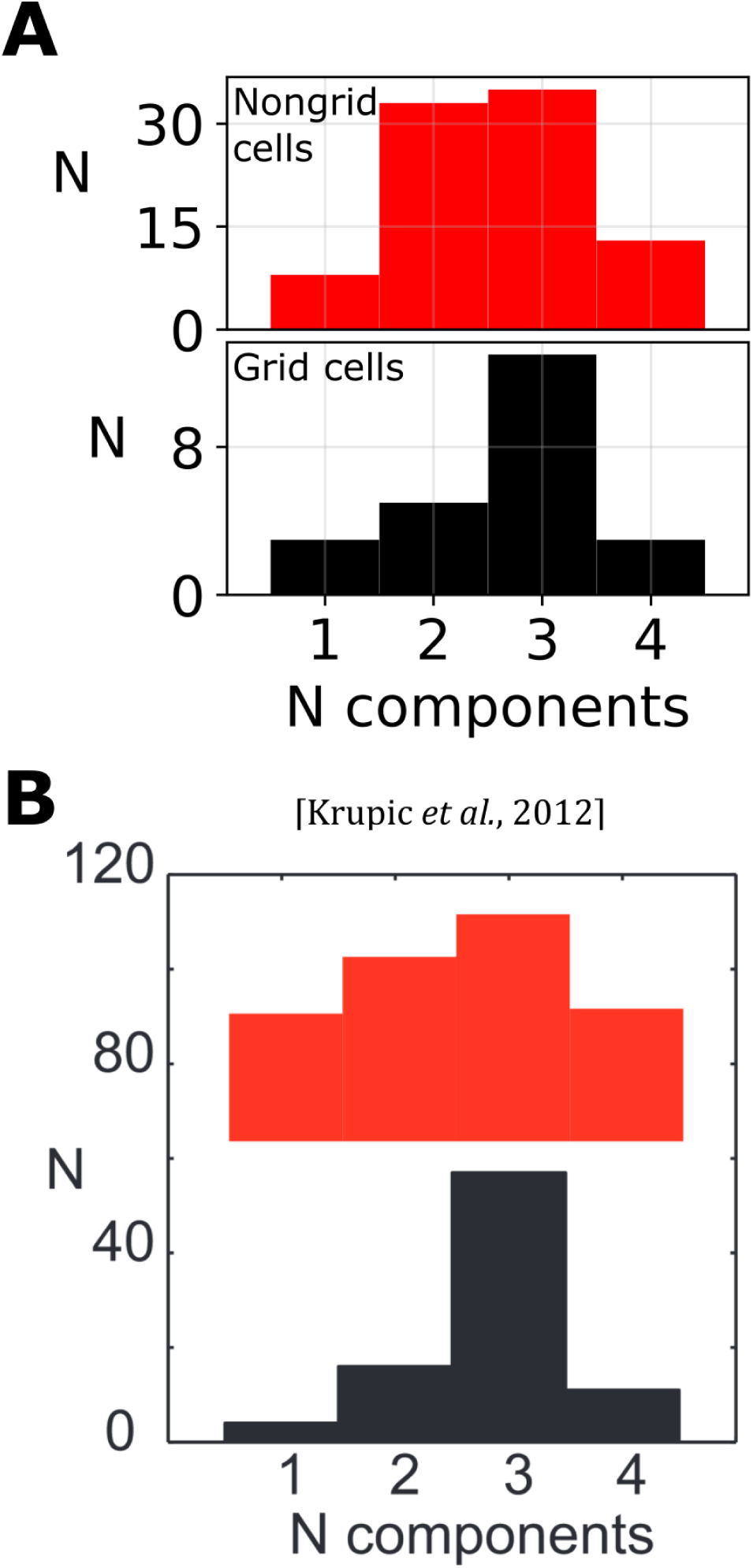
Main Fourier component counts for grid and non-grid cells. Comparison between model and recorded neural main Fourier component counts, where a main component is defined as a peak in the average Fourier spectrogram across angles. (**A**) To obtain the counts, we smooth the polar spectrograms with a Gaussian kernel of width 34 and deviation 26, and we only accept as peaks the local maxima in a range of 40 degrees that are above 0.2 of the maximum value in the spectrogram. (**B**) Main Fourier component counts in the medial entorhinal cortex for grid cells (below, in black) and non-grid cells (above, in red). Taken from (*35*).

**Figure S7:**
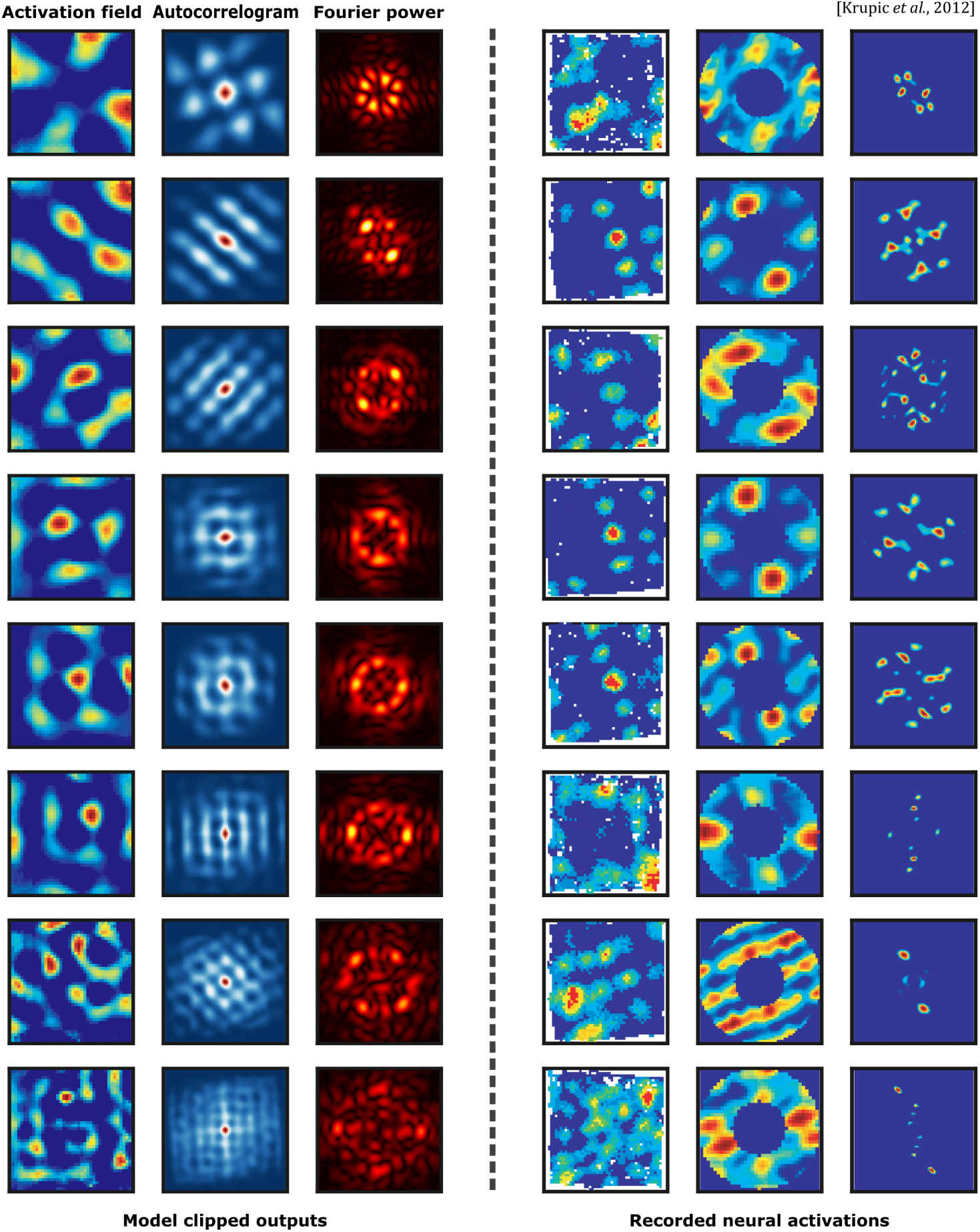
A subset of periodic non-grid cells exhibits square periodicity. Comparison between model features (left columns) and firing patterns observed in rodent studies (right columns). For each model feature, the spatial activation map, autocorrelogram (second-left column), and Fourier spectrogram are shown (third-left column). In contrast with grid cells, the non-grid cells shown here present an autocorrelogram where the central node has 4, instead of 6, peaks at similar distances. Correspondingly, spatial activation maps, both in the case of the model and the neural recordings, contain square-like repeating patterns, as opposed to hexagonal.

**Figure S8:**
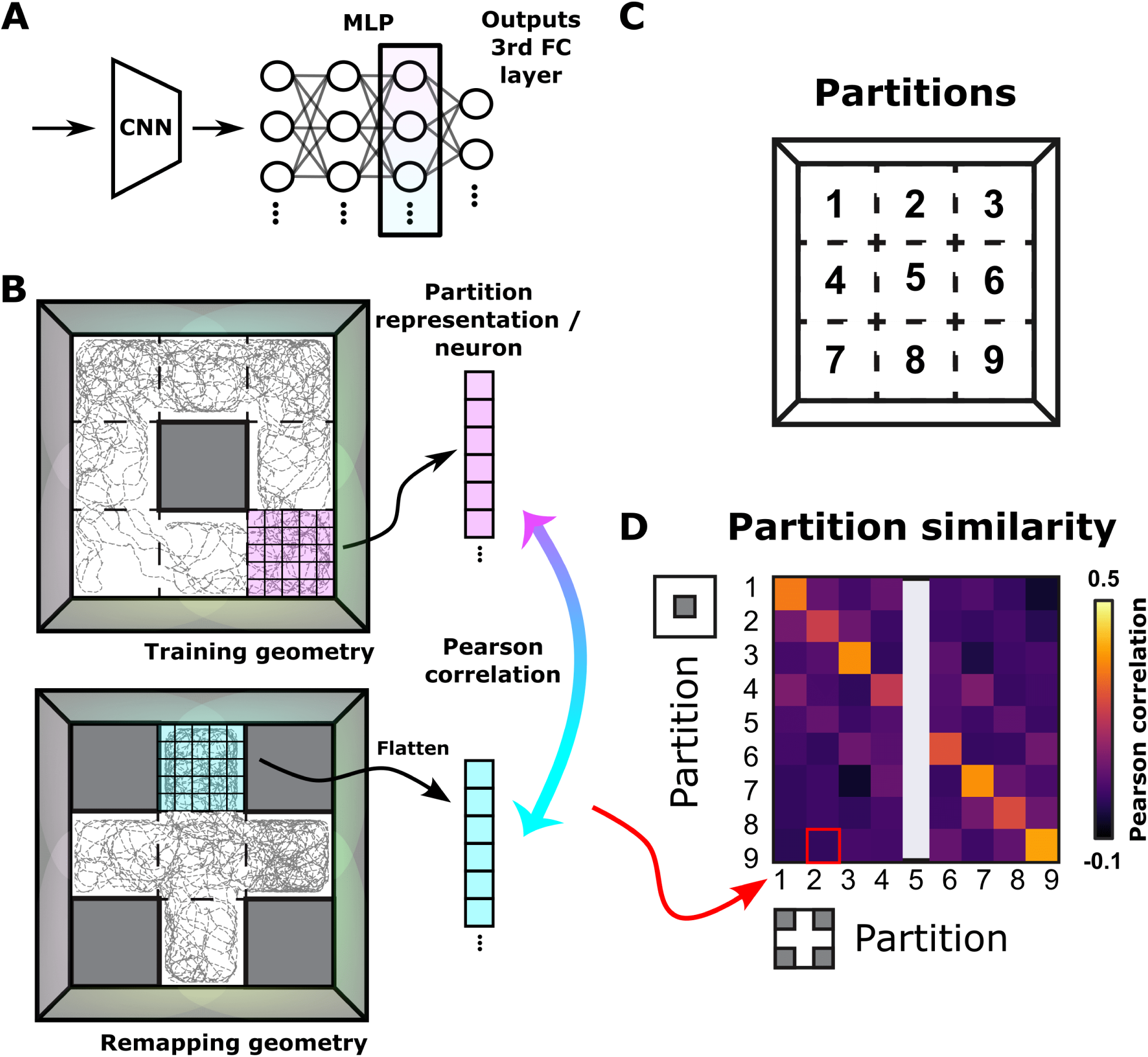
Computing similarity in remapping experiments. (**A**) We evaluate remapping in the second-to-last layer, which resembles place cell activity. (**B**) Partitions in the imagined 3×3 grid are further subdivided into a 5×5 subgrid. The activations of the second-to-last layer are averaged in each of the 25 tiles of the subgrid. The resulting flattened 25-dimensional vector represents each partition, and the similarity is calculated as the Pearson correlation between pairs of partitions from a training to a remapping environment. (**C**) Partition ordering. (**D**) Sub-matrix of the RSM corresponding to the compared training and remapping environments.

**Figure S9:**
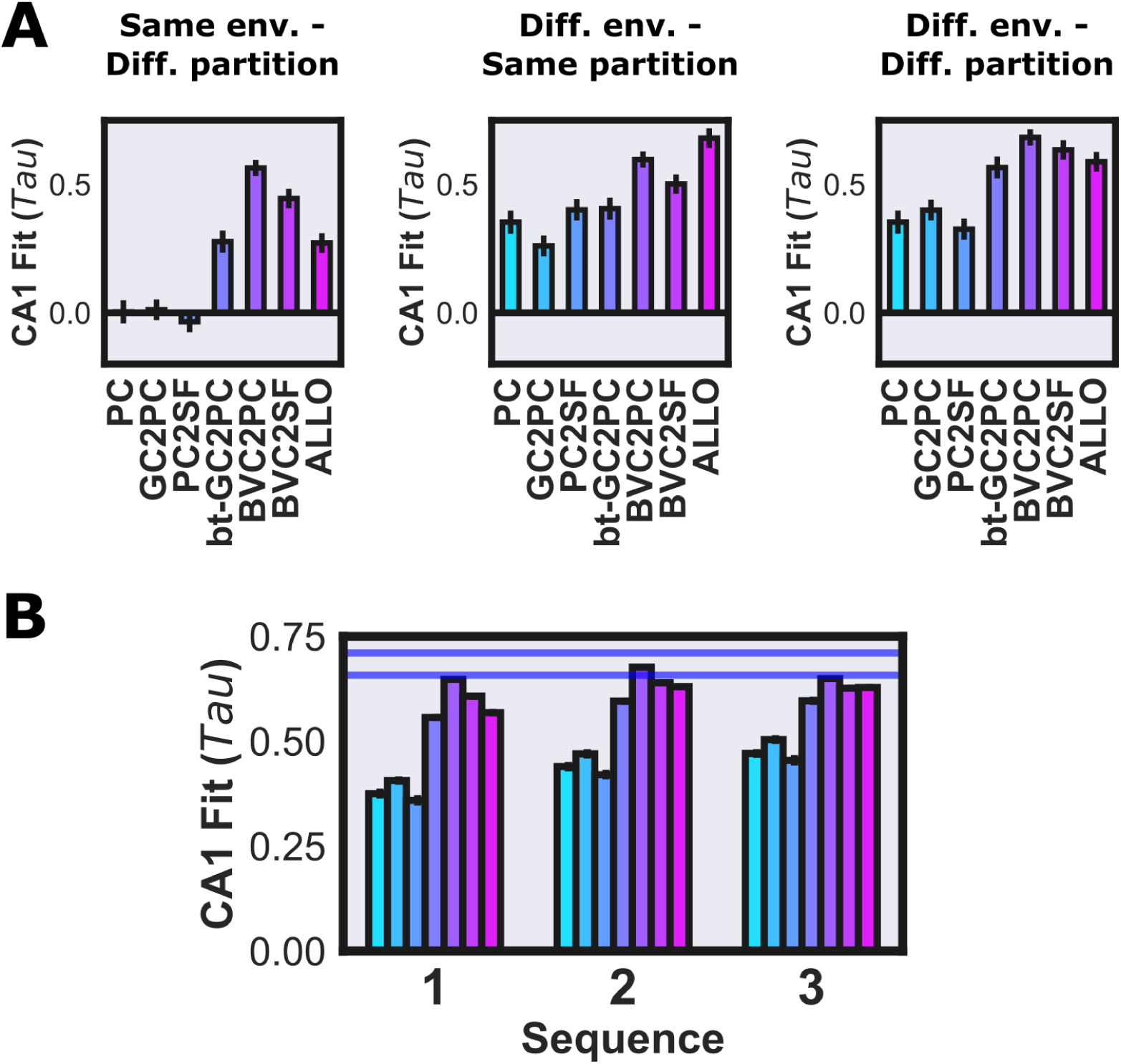
Subset rank-order correlations in remapping experiments. Following (*33*), we evaluate the rank order correlation of RSMs in a subset of the shapes and partitions for our proposed model (ALLO) and compare it with the results in (*33*) for other models. (**A**) We discriminate between three different shapes and partition pairs: same environment and different partition (left), different environment and same partition (center), and both different environment and partition (right). From the corresponding sub-matrices, we calculate rank-order correlations. Remapping is not explained as well in the same environment case, suggesting that hippocampal neurons do remap considerably after leaving and returning to the same environment (something that cannot happen in a static model). (**B**) The 10 different geometries are presented in three repeating sequences spanning 31 days. We discriminated each separate sequence to calculate rank-order correlations, obtaining a convergent performance, similar to the behaviour of biologically-informed models.

**Figure S10:**
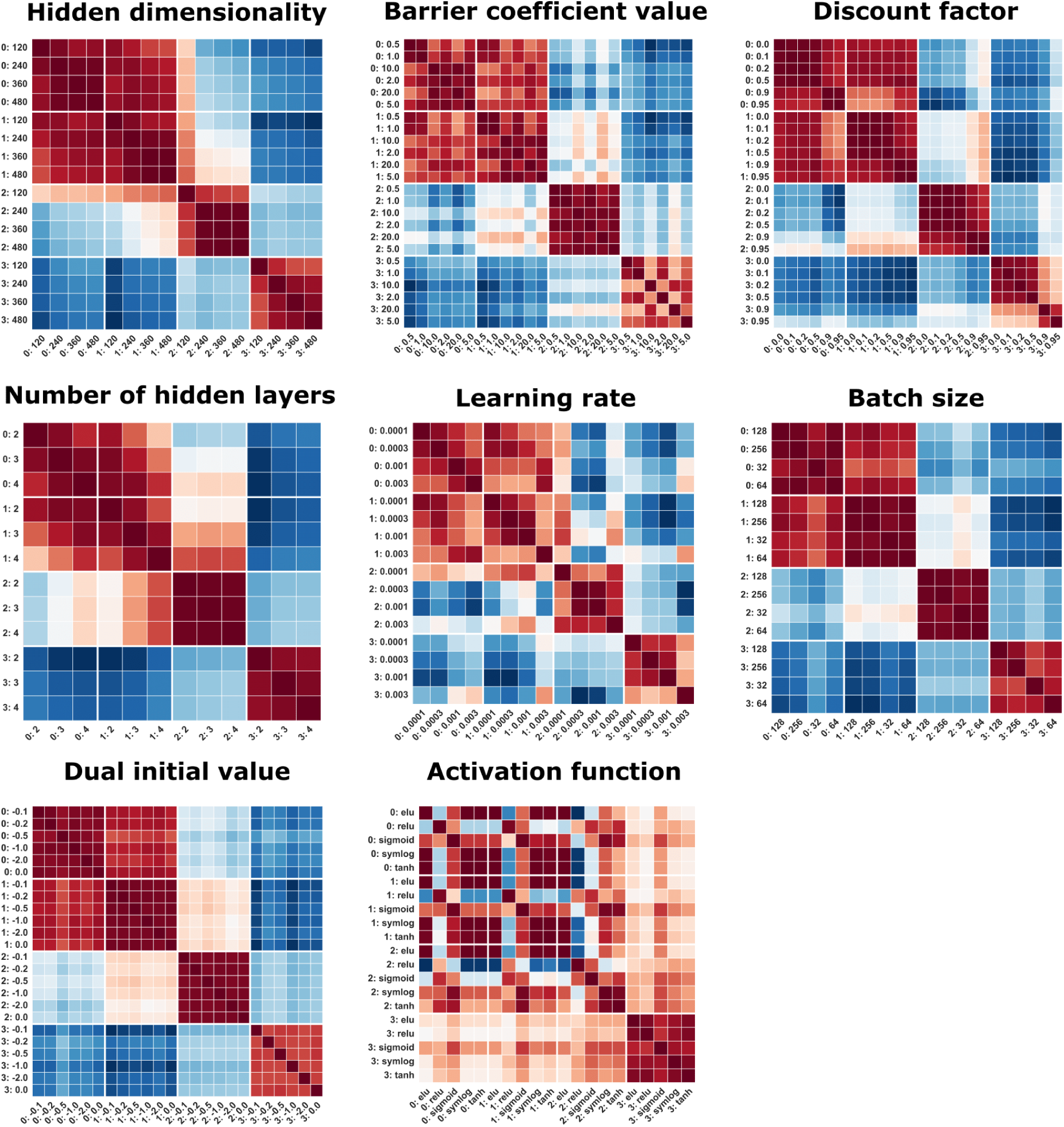
Cosine similarity matrices across remaining hyperparameter values. An entry contains the cosine similarity between two representational similarity analysis (RSM) matrices, each corresponding to a hyperparameter value and a specific layer. Each entry in an RSM reflects similarity between population vectors in corresponding tiles of a 15 × 15 partition of the open arena.

## Notes

### Competing Interest Statement

The authors have declared no competing interest.

